# A revisit to universal single-copy genes in bacterial genomes

**DOI:** 10.1101/2022.04.23.489213

**Authors:** Saidi Wang, Minerva Ventolero, Haiyan Hu, Xiaoman Li

**Affiliations:** Department of Computer Science, University of Central Florida, Orlando, Florida, United States of America; Burnett School of Biomedical Science, College of Medicine, University of Central Florida, Orlando, Florida, United States of America; Genomics and Bioinformatics Cluster, University of Central Florida, Orlando, Florida, United States of America

## Abstract

Universal single-copy genes (USCGs) are widely used for species classification and taxonomic profiling. Despite many studies on USCGs, our understanding of USCGs in bacterial genomes might be out of date, especially how different the USCGs are in different studies, how well a set of USCGs can distinguish two bacterial species, whether USCGs can separate different strains of a bacterial species, to name a few. To fill the void, we studied USCGs in the most updated bacterial RefSeq genomes. We showed that different USCG sets are quite different while coming from highly similar functional categories. We also found that although USCGs occur once in almost all bacterial genomes, each USCG does occur multiple times in certain genomes. We demonstrated that USCGs are reliable markers to distinguish different species while they cannot distinguish different strains of most bacterial species. Our study shed new light on the usage and limitations of USCGs, which will facilitate their applications in evolutionary, phylogenomic, and metagenomic studies.

## Introduction

Universal single-copy genes (USCGs) are marker genes that occur once and only once in almost every genome [1]. Because of this property of ubiquitous existence and uniqueness in each genome, USCGs are widely used to study the evolution and classification of species [2–4]. In the past two decades, with the enormous amount of metagenomic data generated, USCGs are also routinely employed for taxonomic profiling of microbial species and completeness evaluation of metagenome-assembled genomes in shotgun metagenomic studies [5–9].

A widely used marker gene is the 16S rRNA gene, which is thought to be single-copied while shown otherwise and is thus not a perfect USCG [10]. 16S rRNA gene has been used as the gold standard in amplicon sequencing and demonstrated its power for taxonomic profiling in various metagenomic studies [4, 11–14]. It is the most sequenced gene, with its sequences often the only ones we know about a species. Despite its indisputable value and popularity, the 16S rRNA gene may have multiple copies in an unknown genome [10]. Moreover, its sequences might be too conserved to distinguish certain species and/or correctly measure the species divergence [4]. It is thus natural to consider other USCGs in evolution and metagenomics.

Previous studies have identified USCGs other than the 16S rRNA gene [1–4, 9, 15–19]. For instance, Ciccarelli et al. obtained 31 USCGs to reconstruct the tree of life across all three domains [1]. Later, Creevey et al. inferred 40 USCGs from the same set of 191 complete genomes [16], which were then used for taxonomic profiling in metagenomics by the tools mOTUs and mOTUs2 [5, 6]. Wu and Eisen developed the AMPHORA pipeline for automatic phylogenomic analysis of microbial species based on 31 USCGs [4]. These 31 USCGs were combined with 104 archaeal USCGs for phylogenomic studies [19]. Later, the same group developed the phyEco gene sets that comprised 40 USCGs for “all bacteria and archaea” and 114 USCGs for “all bacteria” [3]. Alneberg et al. analyzed all clusters of orthologous groups in 525 genomes and identified 36 USCGs that occurred in >97% of the 525 genomes with a frequency <1.03 per genome [15]. Lan et al. discovered 73 USCGs in >90% of 1897 genomes to classify prokaryotic species [2]. Parks et al. extracted 120 USCGs present in >90% of genomes and single-copied in >95% of their present genomes [9]. These sets of USCGs are inferred from different groups of sequenced genomes with different purposes. They are thus different, although they do share a fraction of USCGs.

With multiple USCGs, in addition to studying evolution, classifying and taxonomic profiling of species, several studies attempted to investigate the strain diversity [7, 20–23]. For instance, Quince et al. developed tools to de novo reconstruct bacterial strain genomes from shotgun metagenomic reads with the aforementioned 36 USCGs [7, 23]. The StrainPhlAn tool infers bacterial strains directly from shotgun reads based on ~200 clade-specific marker genes [22]. Nayfach et al. proposed a pipeline for strain profiling in metagenomic datasets with 15 USCGs [24]. Such studies of bacterial strains are important for understanding drug resistance, microbial diversity, and the cure of various complex diseases [25–30].

Despite many studies on USCGs, our understanding of USCGs is somewhat outdated. For instance, how much agreement is there between the USCG sets from different studies? For a set of USCGs, how universal are these USCGs in the ever-increasing number of sequenced genomes? When defining the above sets of USCGs, to our knowledge, only the first two sets of USCGs were selected by requiring their single-copy occurrence in all 191 genomes. None of the USCG sets is tested on the latest set of RefSeq genomes to show their universalism. Moreover, it is also not clear whether certain USCGs can tell two species apart better than others. In addition, it is unknown whether a set of USCGs such as the 40 USCGs from Creevey et al. [16] contain enough variations to distinguish different strains of a bacterial species.

To address these questions, especially the last one, in this study, we compared different sets of USCGs. We systematically studied how similar a USCG is in different species and strains with the latest set of complete RefSeq genomes. We found that almost every USCG occurs multiple times in certain genomes, while more than 99.4% of USCGs are single-copied in a given genome. We also observed that USCGs together are good for separating species, while they cannot distinguish strains from the same bacterial species in general. Our study provides a more updated picture of USCGs and their potential applications in evolutionary and metagenomic studies.

## Results

### USCG sets are different in gene content while similar in gene function

We compared USCG sets from seven studies [1–4, 9, 15, 16] (Material and Methods, Supplementary Table S1). They were (1). 31 USCGs from Ciccarelli et al.; (2). 31 USCGs from Wu and Eisen; (3). 40 USCGs from Creevey et al.; (4). 40 USCGs from Wu et al. for “all” bacteria and archaea; (5). 36 USCGs from Alneberg et al.; (6). 73 USCGs from Lan et al.; and (7). 120 USCGs from Parks et al. These USCG sets require different universalism across all genomes and uniqueness in individual genomes. For instance, USCGs in the last two sets occurred in only >90% of the genomes, while that was at least >97% in the third to the fifth sets.

We found that even the USCG sets from the same research group could be very different (Figure 1A). For instance, only 18 (77%) of the 31 USCGs from Wu and Eisen were shared with the 40 USCGs from Wu et al. Both sets were inferred by the same research group, with the latter inferred from a much larger number of genomes [3, 4]. Their difference thus implied the significant effect of the genomes used to infer these USCGs On the other hand, the USCG sets from the same research group could be highly consistent as well. For instance, the USCG set from Creevey et al. was a superset of the USCG set from Ciccarelli et al., as the same research group inferred them with the same genomes and a slightly different strategy. However, the additional nine USCGs from Creevey et al. also demonstrated the binding effect of such different strategies on defining USCGs.

**Figure1.**
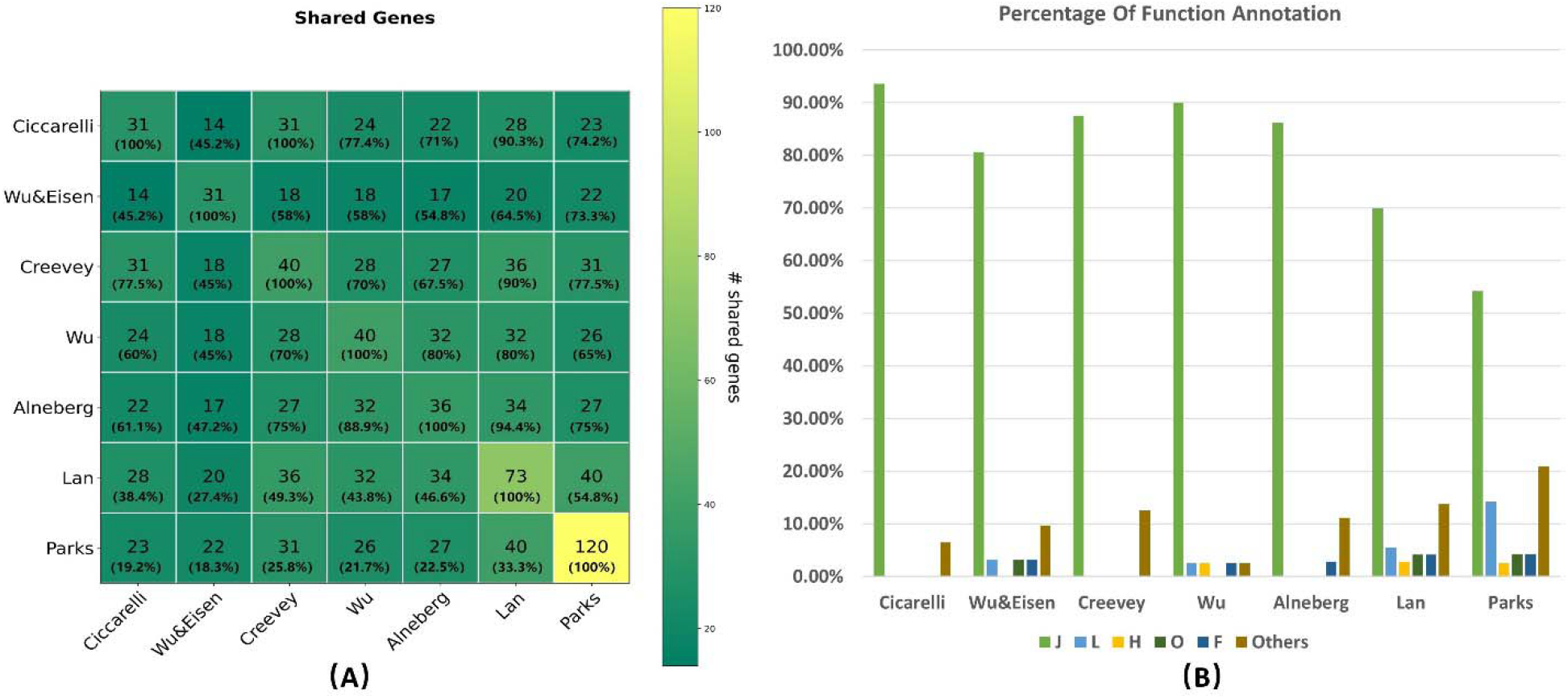
USCG sets and functional categories. (A) Overlap of the seven USCG sets. (B) The percentage of the functional categories of the USCGs in the seven sets.

The USCG sets from different research groups differed more (Supplementary Table S2). For instance, only 14 (45%) of the 31 USCGs from Ciccarelli et al. were shared with those from Wu and Eisen. Again, the difference in these two sets corroborated the effect of different strategies and different genomes. When we considered the two sets refined later by the corresponding research groups, the third and fourth sets, 28 (70%) of the 40 USCGs in these two sets were shared. Although the USCGs in these two sets were still drastically different, they shared much more USCGs than their earlier versions, likely because the number of genomes was large enough to choose a more representative set of genomes by Wu et al. Compared with the sets from Alneberg et al., Lan et al., and Parks et al., the USCGs from Creevey et al. was at least comparable with those from Wu et al., if not better. Overall, the USCG set from Creevey et al. is likely as reliable as any other set, if not more reliable, because of its stricter requirement of universalism and uniqueness, the similar evolutionary trajectory of these USCGs, and likely more representative genomes (consistent results with other sets that used much more genomes).

Despite the difference in the number and members of USCGs, the function of the USCGs in each of the above sets was quite consistent (Figure 1B) [31–35]. By checking the functional annotations at https://www.ncbi.nlm.nih.gov/research/cog, we found that the vast majority of USCGs in every set are annotated with the functional category J (Translation, ribosomal structure, and biogenesis). The remaining USCGs were annotated with other translation and metabolism related categories, such as L (Replication, recombination, and repair), H (Coenzyme transport and metabolism), F (Nucleotide transport and metabolism), O (Posttranslational modification, protein turnover, chaperones), etc. Each category other than J was annotated with much fewer USCGs, from one to a handful of USCGs, compared with several dozen for J. Overall, the USCG sets are enriched with functions related to translation and metabolism, no matter which USCG set is concerned about. For instance, among the 40 USCGs from Wu et al., 35 are involved in the translation process, while the remaining five are related to the cellular metabolic process [3].

### USCGs are indeed universal across species and unique in individual genomes

To our knowledge, the number of genomes used to infer the above USCG sets was no more than 2000 except those from Parks et al. With more than 25,000 complete bacterial genomes at National Center for Biotechnology Information (NCBI), it was unclear whether the USCGs were still universal and unique. We thus studied the occurrence of the USCG sets in the latest set of complete RefSeq genomes (Material and Methods). We focused on the sets from Creevey et al., Wu et al., and Alneberg et al. below because they are widely used in metagenomics and more updated than their previous versions (the first and third set). Moreover, they have more stringent criteria for universalism and uniqueness than the sets from Lan et al. and Parks et al. [2, 9]. In the following, we presented our study on the above three sets.

We found that the 40 USCGs from Creevey et al. were universally distributed in almost each of the 25271 RefSeq genomes (Table 1, Supplementary Table S3, Material and Methods). For every USCG, it occurred in at least 97.7% of the 25271 complete RefSeq genomes. For genomes without a copy of a USCG, we applied BLAST with an arbitrary cutoff of 1E-15 to search for this USCG. BLAST could identify a copy with this cutoff in most of these missed genomes. A tiny fraction of the genomes still did not have a USCG copy, likely due to the cutoff, the quality of the genomes, and the imperfectness of the gene retrieval methods used. Despite these limitations, each USCG occurred in at least 99.7% of the RefSeq genomes. Of note, the number of USCGs occurring in each genome also showed the universal distribution of these USCGs. The mean number of USCGs occurring in each RefSeq genome was 39.6. In other words, almost each of the 40 USCGs occurred in every genome, indicating the universalism of the USCGs from Creevey et al.

**Table 1.** Universalism and uniqueness of USCGs.

|  | mean |  |  | Standard deviation |  |  | minimum |  |  | median |  |  | maximum |  |  |
| --- | --- | --- | --- | --- | --- | --- | --- | --- | --- | --- | --- | --- | --- | --- | --- |
|  | C | W | A | C | W | A | C | W | A | C | W | A | C | W | A |
| fetchMG | 99.0 | 98.6 | 99.0 | 0.5 | 2.7 | 0.8 | 97.7 | 83.9 | 95.9 | 99.3 | 99.3 | 99.3 | 99.6 | 99.7 | 99.6 |
| BLAST | 96.1 | 88.9 | 94.2 | 3.0 | 7.0 | 4.5 | 87.1 | 58.9 | 76.7 | 96.7 | 90.9 | 96.0 | 99.7 | 96.8 | 99.0 |
| universal | 100 | 99.8 | 99.9 | 3.4 | 9.4 | 5.3 | 99.7 | 93.4 | 99.0 | 100 | 99.9 | 100 | 100 | 100 | 100 |
| unique | 98.5 | 97.3 | 98.2 | 1.0 | 3.9 | 2.2 | 95.4 | 82.6 | 89.0 | 99.0 | 99.1 | 99.1 | 99.5 | 99.6 | 99.5 |

The subcolumns with the name C, W, and A refer to the corresponding percentage for the USCGs from Creevey et al., Wu et al., and Alneberg et al., respectively.

We also observed that each of the 40 USCGS from Creevey et al. indeed had only one copy in almost every genome (Table 1). For the 25271 RefSeq genomes, the mean and median percentage of the genomes with only one copy of a USCG were 98.5% and 99.0%, respectively. For individual USCGs, 24 (60%) of the 40 USCGs occurred more than once in <0.2% of the RefSeq genomes, while several other USCGs occurred more than once in >2.0% of the RefSeq genomes. Interestingly, these USCGs that occurred more than once in >2% of the RefSeq genomes were often not identified in other USCG sets. For instance, for the top ten USCGs occurring more than once, only one of them was also identified in the set from Wu et al., suggesting that these USCGs might have occurred multiple times in other studies and were thus filtered by those studies. Overall, the average number of USCGs with only one copy in a genome was 39.4. This average number for the RefSeq genomes suggests that almost all USCGs have only one copy in almost all complete bacterial genomes.

We further studied the universalism and uniqueness of the USCG sets from Wu et al. and Alneberg et al. (Supplementary Table S3). Compared with the set from Creevey et al., the other two sets had a similar median (Figure 2). For instance, the median of the universalism was 99.3% for all three sets, and the corresponding median of the uniqueness was 99.0%, 99.1%, and 99.1%. However, there were more individual USCGs in the other two sets that were not so universal and unique as those from Creevey et al. It is thus evident that the above 40 USCGs from Creevey et al. might be a better USCG set.

**Figure 2.**
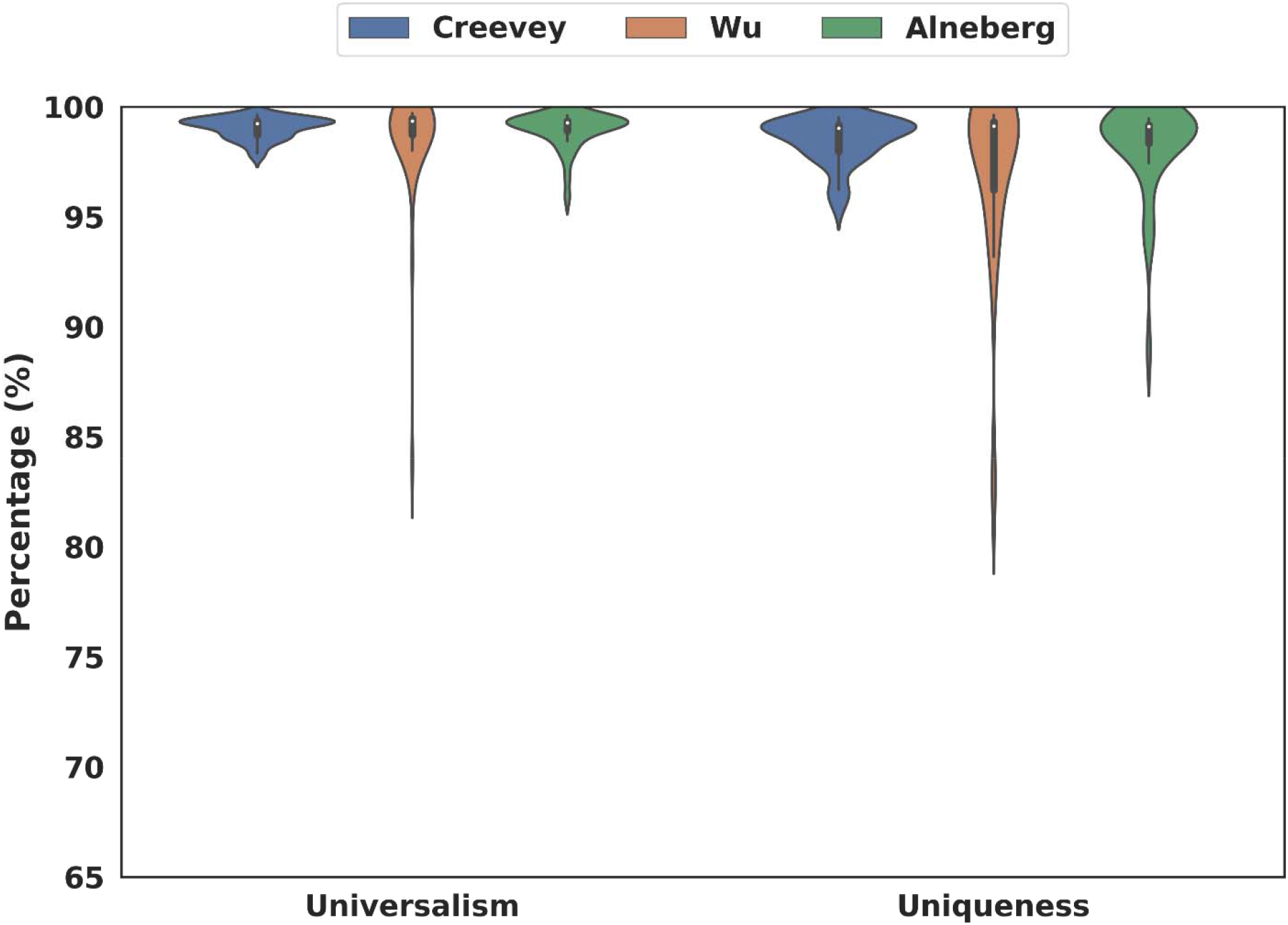
The universalism and uniqueness of the three USCG sets. Comparison of universalism and uniqueness in complete RefSeq genomes.

### USCGs well separate different species, but not strains of most species

With the USCGs indeed universal and unique in bacterial genomes, we investigated how well they together could distinguish two bacterial species in the same genus and whether they together are enough to set two bacterial strains of the same bacterial species apart. To address these questions, we studied all 440 genera with at least two sequenced species genomes and all 747 bacterial species with at least two sequenced strain genomes in the 25271 complete RefSeq genomes (Material and Methods). We found that at least the 40 USCGs from Creevey et al. together were enough to distinguish almost all pairs of species in the same genus. However, they could not separate pairs of strains from most species (Table 2).

**Table 2.** The cumulative distribution of the PID of species pairs and strain pairs.

| PID cutoff |  | <99.5% | <99.6% | <99.7% | <99.8% | <99.9% | <=1 |
| --- | --- | --- | --- | --- | --- | --- | --- |
| Species | Creevey et al. | 97.2 | 97.7 | 98.1 | 98.6 | 99 | 100 |
|  | Wu et al. | 97.2 | 97.7 | 98.2 | 98.5 | 98.8 | 100 |
|  | Alneberg et al. | 97.3 | 97.8 | 98.2 | 98.5 | 98.8 | 100 |
| Strains | Creevey et al. | 7.6 | 8.8 | 10.6 | 15.5 | 25.4 | 100 |
|  | Wu et al. | 5.2 | 6.1 | 7.7 | 20.4 | 43.1 | 100 |
|  | Alneberg et al. | 5.9 | 6.7 | 8.5 | 14.9 | 36.9 | 100 |

The PID cutoff is provided in the first row. The percentage of pairs with their PID smaller than the cutoff is provided in the corresponding entry.

With the Creevey et al. USCG set, we studied the similarity of 40 pairs of corresponding USCGs in every two species from the same genus for all 440 genera with at least two complete genomes (Supplementary Table S4). We measured the similarity by the percentage of identity (PID) in the alignment of the 40 pairs of USCGs from a pair of species because the PID determines how similar the shotgun metagenomics reads from these USCGs are. The PID had a mean and median of 90.2% and 90.6%. However, about 2.3% of species pairs had a PID larger than 99.5%, which was the limit that the current computational tools could distinguish two genomes [7, 36–40]. Given that the total nucleotide length of the 40 USCGs is about 33,852 base pairs, the 99.5% PID means there are at least 170 varied positions in the 40 USCGs required to separate the two species. Therefore, the 40 USCGs could tell most pairs of species apart, which may leave about 2.3% of the pairs of species indistinguishable.

With the same USCG set, we also studied the similarity of 40 pairs of corresponding USCGs in every two strains from the same species for all 747 species with at least two complete genomes (Table 2, Supplementary Table S5). The PID in a pair of strains had a mean of 99.8% and a median of 99.9%. About 91.2% of the PID scores between pairs of strains from the same species were already larger than 99.5% (Table 2). In other words, the 40 USCGs alone may have difficulty in separating more than 91.2% of pairs of bacterial strains.

We also studied how well the USCG set from Wu et al. and from Alneberg et al. separated species and strains (Table 2). Compared with the USCG set from Creevey et al., these two sets had a slightly lower performance, especially in terms of separating strains of the same species. Many more pairs of species and strains had close to 100% PID based on the Wu et al. set compared with the other two sets. Overall, the USCG set from Creevey et al. is likely to perform better in distinguishing species and strains.

## Discussion

We systematically studied the occurrence of USCGs in the most updated list of sequenced bacterial genomes. We showed that USCGs were universally distributed and uniquely present in almost every bacterial species. The 40 USCGs from Creevey et al. together could distinguish different species from the same genus and different strains from the same species well. Barely was there one individual USCG that could do so alone.

There are many other sets of USCGs not compared here. Wrighton et al. used 30 USCGs to measure the completeness of the reconstructed 49 bacterial genomes [18]. Haroon et al. employed the Amphora2 marker genes, including the 31 USCGs for bacteria from Wu and Eisen and 104 archaeal genes, to measure the completeness of the assembled genomes [17]. Rinke et al. estimated the completeness of the assembled genomes based on 139 bacterial and 162□archaeal conserved marker genes [41]. Wu et al. reported 114 USCGs for “all” bacteria in addition to the 40 USCGs compared here [3]. We did not compare these sets of USCGs because they were more for other purposes and not as universal and unique as those compared in this study.

We found that at least three of the seven sets of USCGs were universal and unique. Other sets of USCGs, except the one from Ciccarelli et al., were unlikely to be as universal and unique as the three sets because their selection was not as strict. For instance, Lan et al. and Parks et al. required USCGs to occur in only more than 90% of the genomes instead of all genomes [2, 9]. The Ciccarelli et al. set was already included in the set from Creevey et al.

For the complete RefSeq genomes, we demonstrated that USCGs could distinguish different species but not strains. This was based on the fact that the PID of 99.5% is the limit of current tools to separate similar sequences. When the strain similarity is higher, say more than 99.5%, we are left with only a few dozen variable loci to distinguish different strains. In this case, with the available USCG sets, it may be challenging to distinguish them, even if possible. Note that all aforementioned USCG sets are not created to distinguish bacterial species and strains in microbiomes. In the future, one may hope to generate new USCG sets for this purpose by developing novel methods and integrating different sources of information [42–48]. Alternatively, one may consider overlapping the assembled contigs or metagenome-assembled-genomes with the USCGs so that more polymorphic sites are available to distinguish strains [7, 23].

## Material and Methods

### Seven sets of USCGS

We studied seven sets of USCGs: the 31 set from Ciccarelli et al. [1], the 31 set from Wu and Eisen [4], the 40 set from Creevey et al. [16], the 40 USCGs from Wu et al. [3], the 36 USCGs from Alneberg et al. [15], the 73 USCGs from Lan et al. [2], and the 120 USCGs from Parks et al. [9]. We focused on these sets for the separation of similar strains, not for the completeness of the assembled genomes in metagenomics.

### The complete RefSeq genomes

We studied the complete bacterial Refseq genomes at NCBI. We retrieved the RefSeq genomes from the NCBI website (https://www.ncbi.nlm.nih.gov/genome/browse#!/prokaryotes/) by selecting the ‘Bacteria’ Kingdom and ‘Complete’ Assembly level. There were 25271 genomes with downloadable reference sequences.

### Identification of USCGs in a genome

To identify the corresponding sequences of a USCG in a given genome, we used the tool fetchMG [6]. This tool was used previously to identify and quantify the microbial species in assembled contigs from shotgun metagenomic reads. We used the following command to fetch the USCG sequence in a genome: ./fetchMG.pl -m extraction -p proteins.fasta -o output. Because fetchMG might be imperfect and might miss certain USCG sequences in a genome, we also applied BLAST with the following command to retrieve USCG sequences that were not identified in a genome: psiblast -in_msa marker_aligned.fasta -db db -evalue 1E-15 - num_alignments num_keep. The E-value cutoff used was 1E-15 as previously [3, 4, 19]. We considered the fetched sequences were the corresponding USCG sequences in the genomes. Note that neither fetchMG nor BLAST is perfect for identifying copies of USCGs in a genome, as they sacrifice the accuracy for the sake of speed. However, the identified copies of USCGs are likely to be reliable. We obtained the functional categories of each USCG from https://www.ncbi.nlm.nih.gov/research/cog.

### The USCG similarity measurement

We aligned the extracted USCG protein sequences for every USCG with the tool MAFFT [49]. In other words, we did multiple alignments of the USCG sequences extracted from each genome under consideration for each USCG. In order to assess the similarity of the USCG sequences, we calculated the score based on the blosum62 matrix, the matched number, the mismatch number, the indel number, and the PID. We chose the PID to evaluate the similarity of each pair of USCG sequences because it directly relates to the shotgun metagenomic reads mapped to different USCG sequences. To measure the PID, we obtained the two corresponding sequences in the alignment and removed the loci with an indel versus an indel. We then calculated the PID as the ratio of the number of matched positions to the number of all remaining aligned positions.

## Supporting information

SCG6Li_Supplemental Information

## Acknowledgment

This work was supported by the National Science Foundation (1661414, 2015838, 2120907).

## Author Contributions

H.H. and X.L. conceived the idea. S.W. and M.V. implemented the idea and generated results. S.W., M.V., X.L. and H.H. analyzed the results and wrote the manuscript. All authors reviewed the manuscript.

## Data availability

The NCBI genomes are from https://www.ncbi.nlm.nih.gov/genome/microbes/. The seven sets of USCGs are from the corresponding publications [1–4, 9, 15, 16] and are listed in Table S1.

