## Supplementary material for "A revisit to universal single-copy genes in bacterial genomes": SCG6Li_Supplemental Information

Table S1. USCG sets for each of the seven studies

| Ciccarelli | COG Category | Wu&Eisen | COG Category | Creevey | COG Category | Wu | COG Category | Alneberg | COG Category | Lan | COG Category | Parks | COG Category |
|---|---|---|---|---|---|---|---|---|---|---|---|---|---|
| COG0012 | J | COG0051 | J | COG0012 | J | COG0016 | J | COG0016 | J | COG0006 | E | COG0012 | J |
| COG0016 | J | COG0052 | J | COG0016 | J | COG0048 | J | COG0048 | J | COG0008 | J | COG0013 | J |
| COG0048 | J | COG0080 | J | COG0018 | J | COG0049 | J | COG0049 | J | COG0016 | J | COG0015 | F |
| COG0049 | J | COG0081 | J | COG0048 | J | COG0051 | J | COG0051 | J | COG0018 | J | COG0016 | J |
| COG0052 | J | COG0085 | K | COG0049 | J | COG0052 | J | COG0052 | J | COG0024 | J | COG0018 | J |
| COG0080 | J | COG0087 | J | COG0052 | J | COG0072 | J | COG0060 | J | COG0030 | J | COG0030 | J |
| COG0081 | J | COG0088 | J | COG0080 | J | COG0080 | J | COG0072 | J | COG0048 | J | COG0037 | J |
| COG0087 | J | COG0090 | J | COG0081 | J | COG0081 | J | COG0080 | J | COG0049 | J | COG0049 | J |
| COG0091 | J | COG0092 | J | COG0085 | K | COG0087 | J | COG0081 | J | COG0050 | J | COG0052 | J |
| COG0092 | J | COG0093 | J | COG0087 | J | COG0088 | J | COG0087 | K | COG0051 | J | COG0055 | C |
| COG0093 | J | COG0094 | J | COG0088 | J | COG0089 | J | COG0088 | J | COG0052 | J | COG0060 | J |
| COG0094 | J | COG0097 | J | COG0090 | J | COG0090 | J | COG0089 | J | COG0060 | J | COG0072 | J |
| COG0096 | J | COG0098 | J | COG0091 | J | COG0091 | J | COG0090 | J | COG0072 | J | COG0080 | J |

| | | | | | | | | | | | | | |
|---|---|---|---|---|---|---|---|---|---|---|---|---|---|
| COG0097 | J | COG0099 | J | COG0092 | J | COG0092 | J | COG0091 | J | COG0080 | J | COG0081 | J |
| COG0098 | J | COG0100 | J | COG0093 | J | COG0093 | J | COG0092 | J | COG0081 | J | COG0085 | K |
| COG0099 | J | COG0102 | J | COG0094 | J | COG0094 | J | COG0093 | J | COG0085 | J | COG0086 | K |
| COG0100 | J | COG0103 | J | COG0096 | J | COG0096 | J | COG0094 | J | COG0086 | K | COG0087 | J |
| COG0102 | J | COG0126 | G | COG0097 | J | COG0097 | J | COG0096 | J | COG0087 | J | COG0088 | J |
| COG0103 | J | COG0185 | J | COG0098 | J | COG0098 | J | COG0097 | J | COG0088 | J | COG0090 | J |
| COG0172 | J | COG0195 | K | COG0099 | J | COG0099 | J | COG0100 | J | COG0090 | J | COG0091 | J |
| COG0184 | J | COG0197 | J | COG0100 | J | COG0100 | J | COG0102 | J | COG0091 | J | COG0092 | J |
| COG0186 | J | COG0211 | J | COG0102 | J | COG0102 | J | COG0103 | J | COG0092 | J | COG0096 | J |
| COG0197 | J | COG0222 | J | COG0103 | J | COG0103 | J | COG0130 | J | COG0093 | J | COG0097 | J |
| COG0200 | J | COG0233 | J | COG0124 | J | COG0130 | J | COG0184 | J | COG0094 | J | COG0098 | J |
| COG0201 | U | COG0264 | J | COG0172 | J | COG0150 | F | COG0185 | J | COG0096 | J | COG0100 | J |
| COG0202 | K | COG0290 | J | COG0184 | J | COG0164 | L | COG0186 | J | COG0097 | J | COG0102 | J |
| COG0256 | J | COG0292 | J | COG0185 | J | COG0181 | H | COG0197 | J | COG0098 | J | COG0103 | J |
| COG0495 | J | COG0335 | J | COG0186 | J | COG0184 | J | COG0198 | J | COG0099 | J | COG0124 | J |
| COG0522 | J | COG0358 | L | COG0197 | J | COG0185 | J | COG0200 | J | COG0100 | J | COG0130 | J |

| | | | | | | | | | | | | | |
|---|---|---|---|---|---|---|---|---|---|---|---|---|---|
| COG0525 | J | COG0504 | F | COG0200 | J | COG0186 | J | COG0201 | U | COG0102 | J | COG0143 | J |
| COG0533 | J | COG0691 | O | COG0201 | U | COG0197 | J | COG0244 | J | COG0103 | J | COG0172 | J |
| | | | | COG0202 | K | COG0198 | J | COG0256 | J | COG0112 | E | COG0173 | J |
| | | | | COG0215 | J | COG0200 | J | COG0504 | F | COG0124 | J | COG0187 | L |
| | | | | COG0256 | J | COG0244 | J | COG0532 | J | COG0125 | F | COG0188 | L |
| | | | | COG0495 | J | COG0255 | J | COG0541 | U | COG0130 | J | COG0194 | F |
| | | | | COG0522 | J | COG0256 | J | COG0552 | U | COG0143 | J | COG0195 | K |
| | | | | COG0525 | J | COG0481 | J | | | COG0162 | J | COG0196 | H |
| | | | | COG0533 | J | COG0532 | J | | | COG0172 | J | COG0197 | J |
| | | | | COG0541 | U | COG0533 | J | | | COG0180 | J | COG0198 | J |
| | | | | COG0552 | U | COG0541 | U | | | COG0184 | J | COG0200 | J |
| | | | | | | | | | | COG0185 | J | COG0201 | U |
| | | | | | | | | | | COG0186 | J | COG0202 | K |
| | | | | | | | | | | COG0197 | J | COG0203 | J |
| | | | | | | | | | | COG0198 | J | COG0206 | D |
| | | | | | | | | | | COG0199 | J | COG0215 | J |

| | | | | | | | | | | | | | |
|---|---|---|---|---|---|---|---|---|---|---|---|---|---|
| | | | | | | | | | | COG0201 | U | COG0216 | J |
| | | | | | | | | | | COG0202 | K | COG0223 | J |
| | | | | | | | | | | COG0209 | F | COG0224 | C |
| | | | | | | | | | | COG0231 | J | COG0233 | J |
| | | | | | | | | | | COG0237 | H | COG0244 | J |
| | | | | | | | | | | COG0244 | J | COG0250 | K |
| | | | | | | | | | | COG0250 | K | COG0261 | J |
| | | | | | | | | | | COG0256 | J | COG0264 | J |
| | | | | | | | | | | COG0258 | L | COG0268 | J |
| | | | | | | | | | | COG0361 | J | COG0275 | J |
| | | | | | | | | | | COG0441 | J | COG0290 | J |
| | | | | | | | | | | COG0442 | H | COG0292 | J |
| | | | | | | | | | | COG0459 | O | COG0319 | J |
| | | | | | | | | | | COG0468 | L | COG0322 | L |
| | | | | | | | | | | COG0480 | J | COG0336 | J |
| | | | | | | | | | | COG0492 | O | COG0353 | L |

| | | | | | | | | | | | | | |
|---|---|---|---|---|---|---|---|---|---|---|---|---|---|
| | | | | | | | | | | COG0495 | J | COG0357 | J |
| | | | | | | | | | | COG0504 | F | COG0358 | L |
| | | | | | | | | | | COG0522 | J | COG0359 | J |
| | | | | | | | | | | COG0525 | J | COG0360 | J |
| | | | | | | | | | | COG0526 | O | COG0443 | O |
| | | | | | | | | | | COG0532 | J | COG0468 | L |
| | | | | | | | | | | COG0541 | U | COG0472 | M |
| | | | | | | | | | | COG0550 | L | COG0481 | J |
| | | | | | | | | | | COG0552 | U | COG0482 | J |
| | | | | | | | | | | COG0575 | I | COG0495 | J |
| | | | | | | | | | | COG0592 | L | COG0497 | L |
| | | | | | | | | | | COG1109 | G | COG0504 | F |
| | | | | | | | | | | | | COG0516 | F |
| | | | | | | | | | | | | COG0522 | J |
| | | | | | | | | | | | | COG0528 | F |
| | | | | | | | | | | | | COG0532 | J |

| | | | | | | | | | | | | | |
|---|---|---|---|---|---|---|---|---|---|---|---|---|---|
| | | | | | | | | | | | | COG0533 | J |
| | | | | | | | | | | | | COG0536 | LD |
| | | | | | | | | | | | | COG0539 | J |
| | | | | | | | | | | | | COG0541 | U |
| | | | | | | | | | | | | COG0544 | O |
| | | | | | | | | | | | | COG0552 | U |
| | | | | | | | | | | | | COG0556 | L |
| | | | | | | | | | | | | COG0566 | J |
| | | | | | | | | | | | | COG0571 | K |
| | | | | | | | | | | | | COG0576 | O |
| | | | | | | | | | | | | COG0592 | L |
| | | | | | | | | | | | | COG0593 | L |
| | | | | | | | | | | | | COG0632 | L |
| | | | | | | | | | | | | COG0635 | H |
| | | | | | | | | | | | | COG0653 | U |
| | | | | | | | | | | | | COG0669 | H |

| | | | | | | | | | | | | | |
|---|---|---|---|---|---|---|---|---|---|---|---|---|---|
| | | | | | | | | | | | | COG0690 | U |
| | | | | | | | | | | | | COG0691 | O |
| | | | | | | | | | | | | COG0742 | J |
| | | | | | | | | | | | | COG0749 | L |
| | | | | | | | | | | | | COG0750 | KO |
| | | | | | | | | | | | | COG0771 | M |
| | | | | | | | | | | | | COG0773 | M |
| | | | | | | | | | | | | COG0781 | K |
| | | | | | | | | | | | | COG0799 | J |
| | | | | | | | | | | | | COG0806 | J |
| | | | | | | | | | | | | COG0816 | JKL |
| | | | | | | | | | | | | COG0858 | J |
| | | | | | | | | | | | | COG1066 | L |
| | | | | | | | | | | | | COG1159 | J |
| | | | | | | | | | | | | COG1160 | J |
| | | | | | | | | | | | | COG1185 | J |

| | | | | | | | | | | | | | |
|---|---|---|---|---|---|---|---|---|---|---|---|---|---|
| | | | | | | | | | | | | COG1186 | J |
| | | | | | | | | | | | | COG1197 | KL |
| | | | | | | | | | | | | COG1200 | L |
| | | | | | | | | | | | | COG1214 | J |
| | | | | | | | | | | | | COG1217 | T |
| | | | | | | | | | | | | COG1219 | O |
| | | | | | | | | | | | | COG1314 | U |
| | | | | | | | | | | | | COG1466 | L |
| | | | | | | | | | | | | COG2255 | L |
| | | | | | | | | | | | | COG2812 | L |
| | | | | | | | | | | | | COG4102 | S |

Table S2. Comparison of the USCG sets from the seven studies

| | Ciccarelli | Wu&Eisen | Creevey | Wu | Alneberg | Lan | Parks |
|---|---|---|---|---|---|---|---|
| Ciccarelli | 31/31=100.0% | 14/31=45.2% | 31/31=100.0% | 24/31=77.4% | 22/31=71.0% | 28/31=90.3% | 23/31=74.2% |
| Wu&Eisen | 14/31=45.2% | 31/31=100.0% | 18/31=58.1% | 18/31=58.1% | 17/31=54.8% | 20/31=64.5% | 22/31=71.0% |
| Creevey | 31/40=77.5% | 18/40=45.0% | 40/40=100.0% | 28/40=70.0% | 27/40=67.5% | 36/40=90.0% | 31/40=77.5% |
| Wu | 24/40=60.0% | 18/40=45.0% | 28/40=70.0% | 40/40=100.0% | 32/40=80.0% | 32/40=80.0% | 26/40=65.0% |
| Alneberg | 22/36=61.1% | 17/36=47.2% | 27/36=75.0% | 32/36=88.9% | 36/36=100.0% | 34/36=94.4% | 27/36=75.0% |
| Lan | 28/73=38.4% | 20/73=27.4% | 36/73=49.3% | 32/73=43.8% | 34/73=46.6% | 73/73=100.0% | 40/73=54.8% |
| Parks | 23/120=19.2% | 22/120=18.3% | 31/120=25.8% | 26/120=21.7% | 27/120=22.5% | 40/120=33.3% | 120/120=100.0% |

Table S3. Universalism and uniqueness of USCG sets

| Creevey | | | Wu | | | Alneberg | | |
|---|---|---|---|---|---|---|---|---|
| **USCG** | **Universalism** | **Uniqueness** | **USCG** | **Universalism** | **Uniqueness** | **USCG** | **Universalism** | **Uniqueness** |
| COG0012 | 24950/25271=98.73% | 24890/25271=98.49% | COG0052 | 25106/25271=99.35% | 25024/25271=99.02% | COG0016 | 25017/25271=98.99% | 24966/25271=98.79% |
| COG0016 | 25017/25271=98.99% | 24966/25271=98.79% | COG0051 | 25146/25271=99.51% | 25085/25271=99.26% | COG0048 | 25049/25271=99.12% | 25001/25271=98.93% |
| COG0018 | 24891/25271=98.5% | 24113/25271=95.42% | COG0081 | 25106/25271=99.35% | 25005/25271=98.95% | COG0049 | 25103/25271=99.34% | 25058/25271=99.16% |
| COG0048 | 25049/25271=99.12% | 25001/25271=98.93% | COG0481 | 24951/25271=98.73% | 23928/25271=94.69% | COG0051 | 25146/25271=99.51% | 25049/25271=99.12% |
| COG0049 | 25103/25271=99.34% | 25058/25271=99.16% | COG0532 | 24879/25271=98.45% | 23772/25271=94.07% | COG0052 | 25076/25271=99.23% | 25038/25271=99.08% |
| COG0052 | 25076/25271=99.23% | 25038/25271=99.08% | COG0533 | 24973/25271=98.82% | 23989/25271=94.93% | COG0060 | 24879/25271=98.45% | 23782/25271=94.11% |
| COG0080 | 25115/25271=99.38% | 25046/25271=99.11% | COG0091 | 25116/25271=99.39% | 25082/25271=99.25% | COG0072 | 24944/25271=98.71% | 23914/25271=94.63% |
| COG0081 | 25081/25271=99.25% | 25042/25271=99.09% | COG0541 | 24771/25271=98.02% | 23547/25271=93.18% | COG0080 | 25115/25271=99.38% | 25046/25271=99.11% |
| COG0085 | 24952/25271=98.74% | 24900/25271=98.53% | COG0088 | 25163/25271=99.57% | 25123/25271=99.41% | COG0081 | 25081/25271=99.25% | 25042/25271=99.09% |
| COG0087 | 25086/25271=99.27% | 25047/25271=99.11% | COG0090 | 25134/25271=99.46% | 25114/25271=99.38% | COG0087 | 25086/25271=99.27% | 25047/25271=99.11% |
| COG0088 | 25103/25271=99.34% | 25061/25271=99.17% | COG0103 | 25178/25271=99.63% | 25033/25271=99.06% | COG0088 | 25103/25271=99.34% | 25061/25271=99.17% |
| COG0090 | 25128/25271=99.43% | 25096/25271=99.31% | COG0087 | 25126/25271=99.43% | 25096/25271=99.31% | COG0089 | 25102/25271=99.33% | 25057/25271=99.15% |
| COG0091 | 25088/25271=99.28% | 25056/25271=99.15% | COG0072 | 24944/25271=98.71% | 23937/25271=94.72% | COG0090 | 25128/25271=99.43% | 25096/25271=99.31% |

| | | | | | | | | |
|---|---|---|---|---|---|---|---|---|
| COG0092 | 25103/25271=99.34% | 25064/25271=99.18% | COG0093 | 25113/25271=99.37% | 25041/25271=99.09% | COG0091 | 25088/25271=99.28% | 25056/25271=99.15% |
| COG0093 | 25105/25271=99.34% | 25058/25271=99.16% | COG0098 | 25198/25271=99.71% | 25181/25271=99.64% | COG0092 | 25103/25271=99.34% | 25064/25271=99.18% |
| COG0094 | 25175/25271=99.62% | 25131/25271=99.45% | COG0185 | 25097/25271=99.31% | 25092/25271=99.29% | COG0093 | 25105/25271=99.34% | 25058/25271=99.16% |
| COG0096 | 25138/25271=99.47% | 25097/25271=99.31% | COG0049 | 25115/25271=99.38% | 25090/25271=99.28% | COG0094 | 25175/25271=99.62% | 25131/25271=99.45% |
| COG0097 | 25147/25271=99.51% | 25112/25271=99.37% | COG0197 | 25155/25271=99.54% | 25138/25271=99.47% | COG0096 | 25138/25271=99.47% | 25097/25271=99.31% |
| COG0098 | 25179/25271=99.64% | 25149/25271=99.52% | COG0099 | 24929/25271=98.65% | 24254/25271=95.98% | COG0097 | 25147/25271=99.51% | 25112/25271=99.37% |
| COG0099 | 24877/25271=98.44% | 24843/25271=98.31% | COG0016 | 25065/25271=99.18% | 24169/25271=95.64% | COG0100 | 24923/25271=98.62% | 24888/25271=98.48% |
| COG0100 | 24923/25271=98.62% | 24888/25271=98.48% | COG0200 | 25120/25271=99.4% | 25099/25271=99.32% | COG0102 | 25096/25271=99.31% | 25061/25271=99.17% |
| COG0102 | 25096/25271=99.31% | 25061/25271=99.17% | COG0089 | 25102/25271=99.33% | 25088/25271=99.28% | COG0103 | 25175/25271=99.62% | 25137/25271=99.47% |
| COG0103 | 25175/25271=99.62% | 25137/25271=99.47% | COG0097 | 25181/25271=99.64% | 25160/25271=99.56% | COG0130 | 24477/25271=96.86% | 24317/25271=96.22% |
| COG0124 | 24922/25271=98.62% | 24246/25271=95.94% | COG0080 | 25125/25271=99.42% | 25078/25271=99.24% | COG0184 | 24815/25271=98.2% | 24755/25271=97.96% |
| COG0172 | 25036/25271=99.07% | 24338/25271=96.31% | COG0094 | 25179/25271=99.64% | 25152/25271=99.53% | COG0185 | 25082/25271=99.25% | 25050/25271=99.13% |
| COG0184 | 24815/25271=98.2% | 24755/25271=97.96% | COG0048 | 25072/25271=99.21% | 25056/25271=99.15% | COG0186 | 25101/25271=99.33% | 25062/25271=99.17% |
| COG0185 | 25082/25271=99.25% | 25050/25271=99.13% | COG0255 | 25020/25271=99.01% | 24994/25271=98.9% | COG0197 | 25139/25271=99.48% | 25105/25271=99.34% |
| COG0186 | 25101/25271=99.33% | 25062/25271=99.17% | COG0092 | 25120/25271=99.4% | 24456/25271=96.77% | COG0198 | 25085/25271=99.26% | 25037/25271=99.07% |
| COG0197 | 25139/25271=99.48% | 25105/25271=99.34% | COG0100 | 24932/25271=98.66% | 24898/25271=98.52% | COG0200 | 25109/25271=99.36% | 25075/25271=99.22% |

| | | | | | | | | |
|---|---|---|---|---|---|---|---|---|
| COG0200 | 25109/25271=99.36% | 25075/25271=99.22% | COG0244 | 25125/25271=99.42% | 25106/25271=99.35% | COG0201 | 25035/25271=99.07% | 24624/25271=97.44% |
| COG0201 | 25035/25271=99.07% | 24624/25271=97.44% | COG0096 | 25164/25271=99.58% | 25152/25271=99.53% | COG0244 | 25125/25271=99.42% | 25070/25271=99.2% |
| COG0202 | 25150/25271=99.52% | 24848/25271=98.33% | COG0130 | 24477/25271=96.86% | 24342/25271=96.32% | COG0256 | 25045/25271=99.11% | 25016/25271=98.99% |
| COG0215 | 24843/25271=98.31% | 24320/25271=96.24% | COG0256 | 25071/25271=99.21% | 24957/25271=98.76% | COG0504 | 24234/25271=95.9% | 22484/25271=88.97% |
| COG0256 | 25045/25271=99.11% | 25016/25271=98.99% | COG0184 | 24829/25271=98.25% | 24799/25271=98.13% | COG0532 | 24879/25271=98.45% | 23772/25271=94.07% |
| COG0495 | 24957/25271=98.76% | 24664/25271=97.6% | COG0181 | 21193/25271=83.86% | 20861/25271=82.55% | COG0541 | 24733/25271=97.87% | 24682/25271=97.67% |
| COG0522 | 25137/25271=99.47% | 24775/25271=98.04% | COG0186 | 25152/25271=99.53% | 25122/25271=99.41% | COG0552 | 24700/25271=97.74% | 24666/25271=97.61% |
| COG0525 | 24914/25271=98.59% | 24731/25271=97.86% | COG0102 | 25145/25271=99.5% | 25125/25271=99.42% | | | |
| COG0533 | 24942/25271=98.7% | 24764/25271=97.99% | COG0150 | 23502/25271=93.0% | 21062/25271=83.34% | | | |
| COG0541 | 24733/25271=97.87% | 24682/25271=97.67% | COG0164 | 24527/25271=97.06% | 23100/25271=91.41% | | | |
| COG0552 | 24700/25271=97.74% | 24666/25271=97.61% | COG0198 | 25085/25271=99.26% | 25077/25271=99.23% | | | |

Table S4. PID in 440 genera

| Genus | count | mean | std | min | 25% | 50% | 75% | max |
|---|---|---|---|---|---|---|---|---|
| 1129 | 1485 | 80.1 | 11.2 | 61.2 | 66.8 | 83.6 | 88.9 | 99.7 |
| 83654 | 45 | 98.8 | 0.8 | 97.4 | 98.4 | 98.6 | 99.6 | 100 |
| 89966 | 55 | 88.5 | 3.1 | 83 | 86.3 | 87.1 | 90.7 | 94.8 |
| 133 | 10 | 88 | 3.5 | 85.1 | 85.6 | 86.8 | 89.5 | 96.4 |
| 194 | 1176 | 80 | 6.6 | 69.9 | 76 | 77.4 | 83.9 | 100 |
| 286 | 30135 | 92.3 | 3.8 | 78.6 | 89.3 | 92.4 | 95.5 | 100 |
| 252356 | 10 | 90.7 | 2 | 88.7 | 89.4 | 90.8 | 91 | 95.7 |
| 44249 | 2016 | 82.9 | 4.3 | 75.1 | 79.9 | 82.3 | 84.7 | 99.9 |
| 72763 | 15 | 88 | 4 | 83.5 | 84.8 | 88.3 | 89.1 | 100 |
| 93682 | 3 | 93.9 | 5.3 | 90.9 | 90.9 | 90.9 | 95.5 | 100 |
| 401469 | 3 | 92.9 | 5 | 89.9 | 90 | 90.1 | 94.4 | 98.7 |
| 1386 | 5671 | 84.1 | 7.4 | 61.3 | 78.6 | 81.4 | 91 | 100 |
| 848 | 36 | 86.3 | 7.1 | 80.3 | 81.2 | 81.6 | 95.2 | 99.6 |
| 1301 | 2628 | 89.3 | 3.3 | 74.1 | 87.8 | 88.7 | 90.6 | 100 |
| 1243 | 55 | 92 | 2.6 | 89.7 | 90.3 | 91.2 | 92.4 | 99.9 |
| 75984 | 28 | 94.8 | 4.2 | 87.6 | 94 | 96.6 | 97.1 | 99.8 |
| 28196 | 10 | 94.5 | 1.5 | 92.7 | 93.6 | 93.8 | 95.2 | 98.1 |
| 48736 | 15 | 96.7 | 1.5 | 95.4 | 95.7 | 95.8 | 97.6 | 99.9 |
| 1716 | 2278 | 83 | 3.3 | 77.6 | 80.9 | 82.5 | 83.9 | 99.6 |
| 22 | 1128 | 91.3 | 2.9 | 86.3 | 89.4 | 90.7 | 91.9 | 100 |
| 1678 | 136 | 88.7 | 3.5 | 81.8 | 86 | 89 | 91 | 99.1 |
| 338 | 253 | 95 | 3.1 | 89.5 | 91.6 | 96.7 | 97.5 | 99.8 |
| 33986 | 28 | 91 | 6.9 | 84.6 | 84.9 | 85 | 97.3 | 99.9 |
| 2063 | 6 | 85.4 | 3.7 | 82 | 82.1 | 85.2 | 88.2 | 89.8 |
| 113 | 3 | 91.6 | 4.3 | 88.8 | 89.1 | 89.4 | 93 | 96.5 |

| | | | | | | | | |
|---|---|---|---|---|---|---|---|---|
| 161492 | 1 | 89.1 | nan | 89.1 | 89.1 | 89.1 | 89.1 | 89.1 |
| 390846 | 10 | 94.7 | 1.9 | 93 | 93.3 | 93.5 | 96.6 | 97.9 |
| 943 | 3 | 87.3 | 2.8 | 85.5 | 85.8 | 86 | 88.2 | 90.5 |
| 1839 | 136 | 83.9 | 2.9 | 77.4 | 81.8 | 83.9 | 85.5 | 92 |
| 1827 | 703 | 90.5 | 3.9 | 81.5 | 88.3 | 89.7 | 92.7 | 100 |
| 68287 | 435 | 93.8 | 3.8 | 86.2 | 90.2 | 95.1 | 96.5 | 99.9 |
| 226 | 120 | 94.2 | 2.9 | 89.7 | 93.2 | 94.2 | 95.8 | 100 |
| 167375 | 1 | 90.5 | nan | 90.5 | 90.5 | 90.5 | 90.5 | 90.5 |
| 1763 | 1431 | 90.1 | 2.4 | 83.3 | 88.3 | 89.4 | 91.9 | 99.8 |
| 469 | 1653 | 93.9 | 1.7 | 90 | 92.9 | 93.5 | 94.8 | 100 |
| 1822464 | 153 | 94.3 | 3.2 | 85.5 | 93.4 | 94.8 | 96.5 | 99.5 |
| 620 | 10 | 99.8 | 0.1 | 99.7 | 99.8 | 99.8 | 99.9 | 99.9 |
| 642 | 741 | 97.7 | 2 | 90.4 | 97.7 | 98.1 | 98.4 | 99.8 |
| 482 | 190 | 90.9 | 3.6 | 85.3 | 87.7 | 90.2 | 93.3 | 99 |
| 544 | 2485 | 98.7 | 1.1 | 94.5 | 97.6 | 99.4 | 99.7 | 100 |
| 2651583 | 1 | 97.9 | nan | 97.9 | 97.9 | 97.9 | 97.9 | 97.9 |
| 2093 | 171 | 56.4 | 14.1 | 40.8 | 44.2 | 52.6 | 67.2 | 95.2 |
| 590 | 136 | 99.7 | 0.4 | 98.7 | 99.8 | 99.8 | 99.9 | 100 |
| 2040 | 10 | 87.3 | 4.7 | 83.9 | 84.3 | 84.9 | 90 | 97.7 |
| 1218 | 1 | 74.4 | nan | 74.4 | 74.4 | 74.4 | 74.4 | 74.4 |
| 423349 | 55 | 88.8 | 2.4 | 85.9 | 87.6 | 88.1 | 89.3 | 98.1 |
| 529883 | 3 | 89 | 1.9 | 87.7 | 87.9 | 88.1 | 89.6 | 91.1 |
| 85 | 1 | 79 | nan | 79 | 79 | 79 | 79 | 79 |
| 32008 | 1225 | 94 | 3.3 | 85.4 | 91.1 | 94.6 | 97.2 | 100 |
| 914 | 1 | 76.6 | nan | 76.6 | 76.6 | 76.6 | 76.6 | 76.6 |
| 33882 | 630 | 87.6 | 4.1 | 78 | 86.4 | 87.5 | 89.2 | 99.9 |
| 745 | 10 | 89.6 | 6.1 | 84.8 | 85 | 85.2 | 95.3 | 99.7 |

| | | | | | | | | |
|---|---|---|---|---|---|---|---|---|
| 12916 | 36 | 91.9 | 2.8 | 88.3 | 90 | 91.3 | 95.1 | 97.4 |
| 133925 | 3 | 83.7 | 4.1 | 81 | 81.3 | 81.6 | 85 | 88.4 |
| 33877 | 10 | 91 | 2.3 | 88.5 | 89.2 | 90.9 | 91.8 | 96.1 |
| 613 | 435 | 95.7 | 3.5 | 80.9 | 95.6 | 96.6 | 97.1 | 100 |
| 497 | 78 | 93.4 | 3.8 | 86.4 | 89.6 | 95.1 | 95.6 | 99.8 |
| 2800373 | 6 | 91.8 | 5.4 | 87.1 | 87.1 | 91 | 94.9 | 99.7 |
| 1016 | 105 | 89.7 | 4.7 | 82.3 | 86.8 | 87.6 | 94.3 | 99.3 |
| 662 | 1953 | 92.9 | 2.4 | 87.2 | 91.1 | 92.7 | 94.1 | 100 |
| 2053 | 171 | 90.9 | 3.9 | 86.1 | 87.1 | 90.8 | 93 | 99.8 |
| 57493 | 45 | 87.2 | 5.7 | 81.2 | 82.3 | 84.9 | 92.9 | 99.8 |
| 262 | 120 | 90.2 | 3.7 | 83.5 | 86.7 | 90.8 | 92.3 | 99.8 |
| 1866885 | 703 | 89.9 | 1.8 | 86.2 | 88.9 | 89.8 | 90.6 | 100 |
| 570 | 276 | 97.4 | 2.2 | 88.1 | 97.2 | 97.6 | 99.3 | 100 |
| 122277 | 36 | 99 | 0.4 | 98.3 | 98.7 | 99 | 99.3 | 99.7 |
| 1578 | 351 | 81.7 | 9.6 | 67.2 | 69.9 | 84.6 | 88 | 100 |
| 64895 | 10 | 95.2 | 1 | 94.1 | 94.2 | 95.5 | 95.8 | 97.1 |
| 1279 | 1225 | 89.5 | 3.6 | 83 | 87.6 | 88.8 | 91.6 | 100 |
| 74030 | 21 | 88.8 | 3.7 | 85.8 | 86.8 | 87.4 | 89.1 | 98.6 |
| 670516 | 15 | 97.3 | 0.7 | 96.1 | 96.8 | 97.3 | 97.7 | 99.2 |
| 1654 | 325 | 86.9 | 5.8 | 69.7 | 87.3 | 88.5 | 89.4 | 99.7 |
| 288793 | 3 | 92.7 | 0.6 | 92.1 | 92.4 | 92.7 | 93 | 93.2 |
| 52959 | 120 | 94.1 | 1.7 | 92.3 | 92.9 | 93.5 | 94.2 | 99.9 |
| 445 | 171 | 81.9 | 9.2 | 55.1 | 80.8 | 83 | 87 | 100 |
| 48075 | 3 | 83.2 | 1.3 | 81.7 | 82.8 | 83.9 | 83.9 | 83.9 |
| 363408 | 6 | 93.2 | 6.8 | 86.9 | 87 | 93.2 | 99.4 | 99.5 |
| 12960 | 15 | 89 | 3.3 | 86.4 | 86.6 | 87 | 91.3 | 95.5 |
| 235888 | 15 | 85.3 | 4.1 | 81 | 83.2 | 84.4 | 85.7 | 96.6 |

| | | | | | | | | |
|---|---|---|---|---|---|---|---|---|
| 1883 | 21528 | 90 | 2.9 | 76 | 88.3 | 90.1 | 91.5 | 100 |
| 400634 | 66 | 91.5 | 3.6 | 83.9 | 89.3 | 92.5 | 94 | 97.1 |
| 1649453 | 1 | 91.8 | nan | 91.8 | 91.8 | 91.8 | 91.8 | 91.8 |
| 34072 | 91 | 93.7 | 2.4 | 90.6 | 92.2 | 92.5 | 94.5 | 100 |
| 1142 | 10 | 97.2 | 1.7 | 95.5 | 95.9 | 97.3 | 97.3 | 100 |
| 2745 | 351 | 88.1 | 12.3 | 45.4 | 89 | 90 | 93.8 | 100 |
| 2282523 | 3 | 98.5 | 1.3 | 97.7 | 97.7 | 97.7 | 98.8 | 100 |
| 83614 | 15 | 88.8 | 3.7 | 85.7 | 86.2 | 86.4 | 90.9 | 98.3 |
| 586 | 55 | 96.8 | 1.7 | 94.2 | 95.8 | 96.3 | 98 | 100 |
| 237 | 406 | 88.7 | 2.9 | 84.2 | 86.5 | 87.8 | 89.6 | 98.3 |
| 138 | 6 | 85.9 | 6.6 | 79.4 | 80.2 | 85.3 | 90.6 | 94.3 |
| 1637 | 21 | 90.5 | 5 | 85.3 | 87.4 | 88 | 97.6 | 98.8 |
| 810 | 78 | 84.1 | 5.5 | 78 | 80.7 | 82.2 | 86.1 | 97.7 |
| 1525371 | 1 | 93.2 | nan | 93.2 | 93.2 | 93.2 | 93.2 | 93.2 |
| 107 | 15 | 92.5 | 1.6 | 89.6 | 91.7 | 92.3 | 93 | 96 |
| 629 | 91 | 97.9 | 0.9 | 96 | 97.6 | 98.1 | 98.5 | 100 |
| 1663 | 861 | 88 | 10.8 | 48 | 86.4 | 89.9 | 94 | 100 |
| 1569 | 36 | 87.8 | 4.9 | 82 | 85.2 | 86.4 | 87.4 | 99.9 |
| 93217 | 45 | 95.9 | 3.6 | 88.6 | 96.9 | 97.5 | 97.7 | 99.3 |
| 561 | 15 | 99.6 | 0.1 | 99.5 | 99.5 | 99.7 | 99.7 | 99.9 |
| 1298 | 78 | 87.1 | 3.9 | 81.9 | 83.7 | 87.2 | 87.8 | 99.5 |
| 53246 | 528 | 92.3 | 3.6 | 86.6 | 89.7 | 90.7 | 95.3 | 100 |
| 1485 | 741 | 71.6 | 9.5 | 47.4 | 63.4 | 74.4 | 77.1 | 99.5 |
| 32207 | 28 | 84.4 | 4.3 | 76 | 81.5 | 84.2 | 87.4 | 94.5 |
| 2737 | 21 | 83.5 | 4.5 | 79.8 | 80.9 | 81.3 | 84.8 | 97.1 |
| 990721 | 3 | 75.1 | 16.4 | 65.6 | 65.6 | 65.6 | 79.8 | 94 |
| 357 | 91 | 92.7 | 4.1 | 87.9 | 88.9 | 93.2 | 97.5 | 99.8 |

| | | | | | | | | |
|---|---|---|---|---|---|---|---|---|
| 1372 | 45 | 93.3 | 2.9 | 88.6 | 91.8 | 92.1 | 94.2 | 99 |
| 547 | 1035 | 98.5 | 1.4 | 94.7 | 98.3 | 99.1 | 99.3 | 100 |
| 75 | 28 | 88.8 | 5.1 | 82.6 | 84 | 91.8 | 92.3 | 97.7 |
| 1350 | 210 | 90 | 3.5 | 84.9 | 87.6 | 89.1 | 89.8 | 99.9 |
| 583 | 21 | 98.4 | 0.8 | 97 | 97.7 | 97.8 | 99.2 | 99.7 |
| 2299 | 1 | 89.4 | nan | 89.4 | 89.4 | 89.4 | 89.4 | 89.4 |
| 780 | 78 | 88.6 | 8.9 | 74.3 | 75.8 | 91.5 | 96.4 | 98.8 |
| 2094023 | 1 | 97.8 | nan | 97.8 | 97.8 | 97.8 | 97.8 | 97.8 |
| 234 | 210 | 97.3 | 2.5 | 93.7 | 94.1 | 99.3 | 99.6 | 99.9 |
| 1847 | 28 | 87.6 | 6 | 81.8 | 82.2 | 83.4 | 93.5 | 100 |
| 104267 | 15 | 89.8 | 1.9 | 87.1 | 88.7 | 89.6 | 90.6 | 94.9 |
| 33886 | 45 | 96 | 1.8 | 92.6 | 95.5 | 96.4 | 97 | 99.6 |
| 292691 | 3 | 93 | 1.1 | 92.3 | 92.3 | 92.4 | 93.3 | 94.3 |
| 379 | 1431 | 91.7 | 3.9 | 84.4 | 89 | 89.8 | 94 | 100 |
| 1649486 | 1 | 93.1 | nan | 93.1 | 93.1 | 93.1 | 93.1 | 93.1 |
| 2767358 | 10 | 59.1 | 6.4 | 51.7 | 55.2 | 57 | 63.7 | 71.4 |
| 909656 | 3 | 95 | 3.9 | 92.7 | 92.8 | 92.8 | 96.2 | 99.5 |
| 816 | 276 | 94.4 | 2.1 | 92 | 92.7 | 94 | 95.2 | 100 |
| 119977 | 10 | 90.4 | 8.1 | 81.2 | 81.3 | 94 | 97.4 | 99.9 |
| 1742993 | 15 | 96.4 | 2 | 94.4 | 94.6 | 95.1 | 98.1 | 99.9 |
| 416916 | 10 | 97.4 | 0.9 | 96.6 | 96.9 | 97 | 98.1 | 99.2 |
| 1357 | 78 | 87.9 | 7.6 | 79.3 | 80.1 | 87.8 | 93.8 | 100 |
| 228398 | 3 | 88.3 | 4.9 | 85 | 85.5 | 86.1 | 90 | 93.9 |
| 1125 | 1 | 99 | nan | 99 | 99 | 99 | 99 | 99 |
| 2675229 | 6 | 91.1 | 4.3 | 84.7 | 90.3 | 90.8 | 91.5 | 98.3 |
| 2335 | 6 | 95.8 | 3.6 | 92.4 | 92.5 | 95.7 | 99 | 99.2 |
| 2529408 | 15 | 84.2 | 6.7 | 75.3 | 75.7 | 86.9 | 88.3 | 95.1 |

| | | | | | | | | |
|---|---|---|---|---|---|---|---|---|
| 46239 | 15 | 88.8 | 9.6 | 75.8 | 75.9 | 93.3 | 96.3 | 98 |
| 13687 | 351 | 81.7 | 5.3 | 72.7 | 76.2 | 81.3 | 85.6 | 99 |
| 238749 | 10 | 94 | 4.3 | 90.6 | 90.7 | 90.8 | 98.8 | 99.7 |
| 1041 | 66 | 86.8 | 2.5 | 83.7 | 85.2 | 85.7 | 88.5 | 95 |
| 1434011 | 15 | 96.3 | 1.3 | 95.2 | 95.3 | 96 | 96.3 | 100 |
| 209 | 78 | 71.4 | 5.1 | 64.3 | 68.1 | 71 | 73.1 | 90.8 |
| 374 | 496 | 92 | 2.9 | 86.9 | 90.1 | 90.9 | 94.1 | 100 |
| 336276 | 3 | 96.3 | 2.6 | 94.8 | 94.8 | 94.8 | 97 | 99.3 |
| 1518149 | 6 | 88.3 | 1.7 | 86.7 | 86.8 | 88.2 | 89.3 | 90.7 |
| 203804 | 3 | 62.5 | 0.7 | 61.9 | 62 | 62.2 | 62.8 | 63.3 |
| 28105 | 15 | 92.9 | 3.5 | 89.4 | 90.1 | 90.8 | 95.2 | 99.4 |
| 1813 | 55 | 86.3 | 4.2 | 80 | 83.3 | 85.1 | 88.7 | 96.4 |
| 407 | 36 | 91.2 | 4.9 | 86.5 | 87 | 90.1 | 97.2 | 100 |
| 2675234 | 1 | 82.2 | nan | 82.2 | 82.2 | 82.2 | 82.2 | 82.2 |
| 1707 | 45 | 89.8 | 2.2 | 86.8 | 88.4 | 89.3 | 90.2 | 96.2 |
| 41275 | 153 | 91 | 3.7 | 83.6 | 88.4 | 90 | 93.9 | 99.3 |
| 46255 | 66 | 85.4 | 4.4 | 78.3 | 83.2 | 85 | 87.2 | 97.9 |
| 59732 | 595 | 92.5 | 3.7 | 86.1 | 88.6 | 94.1 | 94.9 | 100 |
| 53457 | 3 | 89.2 | 3.9 | 86.5 | 86.9 | 87.3 | 90.5 | 93.7 |
| 44000 | 1 | 93.4 | nan | 93.4 | 93.4 | 93.4 | 93.4 | 93.4 |
| 165696 | 28 | 87.9 | 2.8 | 84.3 | 85.7 | 86.9 | 90.5 | 93.3 |
| 165695 | 136 | 92.1 | 3.4 | 84.6 | 91.1 | 92.6 | 93.7 | 100 |
| 407217 | 6 | 97.3 | 1.1 | 95.6 | 96.8 | 97.5 | 97.7 | 98.8 |
| 447792 | 3 | 99.5 | 0.3 | 99.3 | 99.3 | 99.3 | 99.6 | 99.9 |
| 838 | 78 | 88.2 | 3.9 | 83.7 | 85.2 | 86.6 | 90.1 | 98.6 |
| 1375 | 15 | 79.1 | 5.7 | 75.3 | 76.4 | 77.8 | 78.8 | 98.6 |
| 2742 | 66 | 92.2 | 3.3 | 84.6 | 90 | 92.8 | 94 | 100 |

| | | | | | | | | |
|---|---|---|---|---|---|---|---|---|
| 579 | 3 | 95.8 | 2.6 | 94.3 | 94.3 | 94.3 | 96.5 | 98.8 |
| 773 | 153 | 87.2 | 5 | 77.7 | 84.9 | 87.1 | 88.3 | 99.9 |
| 129337 | 153 | 96.5 | 2.3 | 92.1 | 96.2 | 96.8 | 97.9 | 99.9 |
| 29465 | 15 | 97.5 | 1.3 | 95.5 | 96.4 | 97 | 98.5 | 99.6 |
| 28065 | 28 | 87.2 | 1.4 | 85.1 | 86.1 | 87 | 88.2 | 90.8 |
| 400060 | 3 | 86.7 | 1 | 86.1 | 86.1 | 86.1 | 87 | 87.9 |
| 748770 | 3 | 97.3 | 1.5 | 96.4 | 96.5 | 96.5 | 97.8 | 99 |
| 106589 | 55 | 94.7 | 1.9 | 91.4 | 93.5 | 94.4 | 95.1 | 100 |
| 958 | 10 | 85.2 | 9.1 | 74.8 | 75.4 | 89.2 | 91.2 | 99.4 |
| 106633 | 1 | 93.2 | nan | 93.2 | 93.2 | 93.2 | 93.2 | 93.2 |
| 2132 | 171 | 70.7 | 9.3 | 56.4 | 64.5 | 65.5 | 78 | 100 |
| 893 | 1 | 71.8 | nan | 71.8 | 71.8 | 71.8 | 71.8 | 71.8 |
| 33926 | 21 | 75.1 | 16.7 | 58.5 | 59.4 | 64.2 | 98.5 | 99.4 |
| 58050 | 15 | 97.9 | 1.2 | 96 | 96.6 | 98 | 98.8 | 99.8 |
| 953 | 276 | 88.9 | 5.1 | 80.7 | 85 | 88.5 | 90.1 | 100 |
| 626 | 10 | 96.2 | 0.5 | 95.7 | 95.9 | 96 | 96.6 | 97 |
| 713 | 3 | 93.5 | 1.6 | 92.2 | 92.6 | 93 | 94.1 | 95.2 |
| 44013 | 378 | 92.4 | 4.9 | 81.7 | 90.5 | 94.4 | 95.9 | 99.3 |
| 535 | 28 | 95.3 | 2.3 | 93 | 93.3 | 93.5 | 97.1 | 99.8 |
| 2803850 | 10 | 90.3 | 5.8 | 83.3 | 83.7 | 94 | 94.5 | 96.4 |
| 1912216 | 6 | 91.7 | 4.5 | 87.2 | 87.8 | 91.5 | 95.5 | 96.5 |
| 302485 | 10 | 97.4 | 1.3 | 95.6 | 96 | 98.2 | 98.3 | 98.8 |
| 2767885 | 3 | 96.4 | 0.9 | 95.6 | 95.9 | 96.3 | 96.8 | 97.4 |
| 776 | 3 | 79 | 2.3 | 77 | 77.7 | 78.4 | 80 | 81.5 |
| 53945 | 1 | 86 | nan | 86 | 86 | 86 | 86 | 86 |
| 2321111 | 21 | 89.8 | 1.8 | 86.7 | 88.8 | 89.9 | 90.9 | 95 |
| 59739 | 6 | 94.4 | 3.7 | 91.7 | 91.9 | 92.2 | 97 | 99.7 |

| | | | | | | | | |
|---|---|---|---|---|---|---|---|---|
| 165697 | 45 | 92.8 | 1.7 | 90.3 | 91.6 | 92.7 | 93.7 | 99.2 |
| 551 | 28 | 90.6 | 9.1 | 74.4 | 88.6 | 95.5 | 96.4 | 98.7 |
| 2767886 | 1 | 91.4 | nan | 91.4 | 91.4 | 91.4 | 91.4 | 91.4 |
| 1506553 | 3 | 83.2 | 6 | 79.6 | 79.8 | 79.9 | 85.1 | 90.2 |
| 1030 | 3 | 92.3 | 2.6 | 90.6 | 90.8 | 91.1 | 93.2 | 95.3 |
| 2675233 | 10 | 90.3 | 3.9 | 85.9 | 86.9 | 89.1 | 94.2 | 95.7 |
| 724 | 21 | 91.9 | 4.7 | 85.2 | 87.7 | 91.1 | 96.3 | 98.3 |
| 1696 | 78 | 77.3 | 19.7 | 46.5 | 48 | 81.3 | 92.1 | 98.6 |
| 97050 | 6 | 90.5 | 3.1 | 87.9 | 88.8 | 89.7 | 90.6 | 96.5 |
| 1827195 | 3 | 95.1 | 4.1 | 92.7 | 92.8 | 92.8 | 96.3 | 99.8 |
| 239759 | 10 | 84.3 | 4.8 | 80.4 | 80.8 | 81 | 88.3 | 93.3 |
| 157 | 21 | 75.9 | 7.3 | 69.4 | 70.4 | 77.2 | 77.8 | 98.6 |
| 83618 | 6 | 89.7 | 1.9 | 88 | 88.6 | 89.3 | 90 | 93.2 |
| 84111 | 3 | 91.9 | 4 | 89.5 | 89.6 | 89.7 | 93.1 | 96.5 |
| 68 | 120 | 87 | 4.4 | 79.5 | 85 | 86.1 | 88.5 | 100 |
| 65496 | 10 | 89.8 | 8.5 | 79.7 | 80.2 | 94.2 | 96.6 | 99.9 |
| 60136 | 66 | 89.7 | 3.3 | 86 | 87 | 89.3 | 90.6 | 99.1 |
| 352 | 1 | 97.4 | nan | 97.4 | 97.4 | 97.4 | 97.4 | 97.4 |
| 59734 | 3 | 97 | 0.2 | 96.8 | 96.9 | 97 | 97.1 | 97.2 |
| 1873 | 36 | 89.9 | 3.6 | 80.5 | 87.7 | 90 | 92.5 | 96.6 |
| 34008 | 3 | 91.3 | 4.5 | 88.5 | 88.7 | 88.9 | 92.7 | 96.4 |
| 84567 | 28 | 91.9 | 3.3 | 88.8 | 89.3 | 89.7 | 93.9 | 98.9 |
| 69578 | 3 | 98.1 | 0.9 | 97.5 | 97.5 | 97.6 | 98.3 | 99.1 |
| 1378 | 15 | 92.4 | 4.2 | 86.9 | 87.3 | 93.6 | 94.8 | 99.9 |
| 125287 | 3 | 83.7 | 3.7 | 81.5 | 81.5 | 81.6 | 84.8 | 88 |
| 478070 | 28 | 95.6 | 3.2 | 92.6 | 93.1 | 94.3 | 99.6 | 100 |
| 202746 | 55 | 88.6 | 3.4 | 86 | 86.7 | 87.4 | 88.9 | 99.7 |

| | | | | | | | | |
|---|---|---|---|---|---|---|---|---|
| 2767879 | 21 | 92.4 | 3.4 | 88.5 | 89.3 | 90.3 | 95.2 | 98.8 |
| 2420 | 1 | 90.8 | nan | 90.8 | 90.8 | 90.8 | 90.8 | 90.8 |
| 46913 | 21 | 86.5 | 3.4 | 82 | 83.7 | 85.1 | 88.5 | 94.7 |
| 204037 | 28 | 97.3 | 1.1 | 95.2 | 96.6 | 97.7 | 97.9 | 99.9 |
| 69965 | 15 | 93.3 | 4.6 | 89.1 | 89.2 | 89.3 | 97.6 | 99.8 |
| 160674 | 10 | 94.7 | 5.4 | 88.2 | 88.6 | 97.5 | 99.4 | 100 |
| 1177 | 120 | 92.3 | 2.7 | 89.3 | 90.4 | 91.2 | 93 | 99.3 |
| 283 | 45 | 88.7 | 4.4 | 82.4 | 85.4 | 86.4 | 92.2 | 97.1 |
| 141948 | 6 | 92.2 | 2 | 90.3 | 90.8 | 91.4 | 93.8 | 95 |
| 261825 | 1 | 98.9 | nan | 98.9 | 98.9 | 98.9 | 98.9 | 98.9 |
| 57665 | 15 | 95.4 | 3.3 | 91.2 | 91.8 | 94.9 | 98.6 | 100 |
| 40544 | 3 | 89.5 | 2.8 | 87.7 | 87.9 | 88.1 | 90.4 | 92.8 |
| 76831 | 21 | 95 | 3.1 | 91.2 | 92.7 | 93.3 | 98.2 | 99.9 |
| 47251 | 6 | 74.6 | 2.5 | 71.4 | 74 | 74.1 | 75.1 | 78.9 |
| 517 | 105 | 92.1 | 2.8 | 87.5 | 90.3 | 92.1 | 92.9 | 99.6 |
| 34098 | 21 | 83.2 | 7.6 | 75.8 | 77.7 | 78.7 | 92 | 96.9 |
| 33057 | 6 | 90.5 | 3.6 | 87 | 87.4 | 90.5 | 93.6 | 94.2 |
| 1810868 | 3 | 94 | 2 | 92.7 | 92.8 | 92.9 | 94.6 | 96.3 |
| 55968 | 15 | 90.2 | 3.3 | 85.7 | 86.8 | 90.1 | 92 | 96.5 |
| 104264 | 6 | 91.9 | 4.8 | 87.8 | 88.2 | 91 | 93.8 | 99.9 |
| 146785 | 1 | 98.9 | nan | 98.9 | 98.9 | 98.9 | 98.9 | 98.9 |
| 2013 | 6 | 93.2 | 2.7 | 90.1 | 91.7 | 92.8 | 94.4 | 97.6 |
| 963 | 10 | 95.7 | 2.7 | 92.6 | 92.8 | 96.3 | 98 | 98.9 |
| 2767893 | 10 | 90.7 | 3.6 | 86.6 | 87.3 | 90.9 | 93.5 | 96.4 |
| 191028 | 1 | 96 | nan | 96 | 96 | 96 | 96 | 96 |
| 444051 | 6 | 98.3 | 1.9 | 96.6 | 96.6 | 98.3 | 100 | 100 |
| 265 | 105 | 88.6 | 3.1 | 84.1 | 86.7 | 87.6 | 89.3 | 99.4 |

| | | | | | | | | |
|---|---|---|---|---|---|---|---|---|
| 2742598 | 21 | 88.8 | 3.2 | 85.1 | 86.3 | 87.5 | 91.9 | 94.4 |
| 59753 | 3 | 88.2 | 1 | 87.4 | 87.6 | 87.8 | 88.5 | 89.3 |
| 28231 | 3 | 81.7 | 5.8 | 78.1 | 78.3 | 78.6 | 83.5 | 88.4 |
| 84406 | 21 | 86.9 | 6.1 | 79.7 | 80.7 | 87 | 93.3 | 98.3 |
| 2923352 | 6 | 65.2 | 3.9 | 59.7 | 63.2 | 64.8 | 67.7 | 70.5 |
| 673534 | 3 | 84.8 | 8.2 | 79.3 | 80.1 | 80.9 | 87.6 | 94.2 |
| 434 | 36 | 91.6 | 5.5 | 83 | 88.4 | 91 | 98.9 | 99.8 |
| 2759736 | 6 | 97.1 | 1.3 | 96 | 96.2 | 96.8 | 97.3 | 99.4 |
| 2433 | 1 | 90.6 | nan | 90.6 | 90.6 | 90.6 | 90.6 | 90.6 |
| 622681 | 15 | 88.3 | 3.9 | 84.5 | 85.7 | 86.3 | 89.8 | 96.8 |
| 53335 | 153 | 93.2 | 5.8 | 76.2 | 93.3 | 94.7 | 96.2 | 100 |
| 875170 | 1 | 92 | nan | 92 | 92 | 92 | 92 | 92 |
| 32067 | 21 | 91.3 | 5 | 85 | 85.5 | 90.6 | 96.1 | 98.1 |
| 40323 | 171 | 94.5 | 2.9 | 90.7 | 91.9 | 93.6 | 97.2 | 99.9 |
| 48073 | 45 | 91.8 | 2.4 | 88.1 | 90.3 | 91 | 94.5 | 96.8 |
| 125216 | 10 | 88.8 | 7.4 | 83 | 83.1 | 83.5 | 96.7 | 99.6 |
| 1769012 | 3 | 82.7 | 3.7 | 80.3 | 80.5 | 80.8 | 83.8 | 86.9 |
| 836 | 10 | 76.6 | 2.3 | 73.4 | 75.2 | 76.3 | 77.6 | 80.3 |
| 441 | 6 | 94.9 | 3.2 | 92.8 | 92.8 | 92.8 | 96.9 | 99.7 |
| 246875 | 3 | 90.6 | 1.1 | 89.4 | 90.1 | 90.8 | 91.2 | 91.6 |
| 186650 | 3 | 93.6 | 2.5 | 92.1 | 92.2 | 92.3 | 94.4 | 96.5 |
| 1835 | 6 | 94 | 3.7 | 89.5 | 91.3 | 93.8 | 96.8 | 98.4 |
| 1855416 | 1 | 84.7 | nan | 84.7 | 84.7 | 84.7 | 84.7 | 84.7 |
| 1817 | 78 | 87.8 | 3.7 | 77.9 | 86 | 88.7 | 90.1 | 96.7 |
| 906 | 3 | 90.5 | 1 | 89.7 | 90 | 90.3 | 90.9 | 91.6 |
| 149698 | 78 | 90.9 | 2.8 | 86.6 | 89.5 | 89.8 | 92.4 | 99 |
| 1069494 | 3 | 89.8 | 3.6 | 87.6 | 87.7 | 87.8 | 90.8 | 93.9 |

| | | | | | | | | |
|---|---|---|---|---|---|---|---|---|
| 55080 | 36 | 84.3 | 8.2 | 70.9 | 82 | 84.2 | 91.3 | 99.9 |
| 52972 | 6 | 92.8 | 3.9 | 89.9 | 90 | 91.8 | 93.4 | 100 |
| 2767888 | 3 | 89.2 | 6.1 | 85.6 | 85.7 | 85.7 | 91 | 96.3 |
| 135575 | 6 | 93.9 | 3.2 | 91.3 | 91.5 | 93.2 | 94.9 | 99.4 |
| 329857 | 6 | 88.4 | 2.9 | 85.4 | 86 | 87.9 | 91 | 92 |
| 102106 | 1 | 83.2 | nan | 83.2 | 83.2 | 83.2 | 83.2 | 83.2 |
| 1865 | 21 | 88.2 | 4.5 | 81.9 | 85.6 | 88.5 | 89.2 | 100 |
| 2035811 | 6 | 88.6 | 2.9 | 85.6 | 85.9 | 88.8 | 91.2 | 91.6 |
| 1234 | 21 | 81.8 | 3.4 | 78.1 | 79 | 81.3 | 83.5 | 90.6 |
| 1186 | 28 | 86.4 | 2.6 | 83.9 | 85 | 85.5 | 86.9 | 96.1 |
| 416 | 15 | 89.2 | 3.4 | 84.7 | 87.1 | 88.3 | 90.8 | 98.9 |
| 80865 | 10 | 98.5 | 1.2 | 96.8 | 97.5 | 99 | 99.1 | 99.9 |
| 361177 | 3 | 86.2 | 2.9 | 84 | 84.5 | 85.1 | 87.3 | 89.5 |
| 269258 | 15 | 91.8 | 4.4 | 88.3 | 89.2 | 89.4 | 94 | 100 |
| 31988 | 6 | 98.1 | 1.7 | 96.4 | 96.5 | 98.1 | 99.6 | 99.8 |
| 1263 | 1 | 68.5 | nan | 68.5 | 68.5 | 68.5 | 68.5 | 68.5 |
| 2425 | 1 | 90.2 | nan | 90.2 | 90.2 | 90.2 | 90.2 | 90.2 |
| 270 | 6 | 92.3 | 2.4 | 90.4 | 90.5 | 91.5 | 92.8 | 96.7 |
| 504090 | 1 | 86.9 | nan | 86.9 | 86.9 | 86.9 | 86.9 | 86.9 |
| 308865 | 28 | 96.6 | 3.6 | 90.5 | 95.4 | 98.8 | 99.1 | 99.8 |
| 34037 | 3 | 98.9 | 0.9 | 98.4 | 98.4 | 98.4 | 99.2 | 99.9 |
| 1060 | 3 | 84.6 | 1.9 | 83.2 | 83.5 | 83.8 | 85.3 | 86.8 |
| 28895 | 1 | 94.9 | nan | 94.9 | 94.9 | 94.9 | 94.9 | 94.9 |
| 2321114 | 6 | 91.2 | 5 | 86.8 | 87 | 90.5 | 94 | 98.7 |
| 29580 | 36 | 88.4 | 12.7 | 65.7 | 84.2 | 96.3 | 98.5 | 99.5 |
| 2147 | 3 | 81.6 | 1.7 | 79.9 | 80.8 | 81.7 | 82.5 | 83.3 |
| 28253 | 10 | 93.5 | 3.7 | 89 | 89.5 | 95.3 | 96.3 | 98.3 |

| | | | | | | | | |
|---|---|---|---|---|---|---|---|---|
| 413 | 6 | 85.9 | 9.8 | 76.7 | 77 | 85.3 | 94.7 | 95.7 |
| 657 | 3 | 92.9 | 1 | 91.7 | 92.6 | 93.4 | 93.5 | 93.5 |
| 290174 | 6 | 92.2 | 2.3 | 90.5 | 90.8 | 91.5 | 92.5 | 96.5 |
| 1778601 | 10 | 95.5 | 5.2 | 89.5 | 89.5 | 99.5 | 99.6 | 99.7 |
| 153265 | 3 | 94.9 | 4.4 | 92.3 | 92.3 | 92.3 | 96.2 | 100 |
| 2747 | 10 | 87 | 4.5 | 83.8 | 84.4 | 85.3 | 87.2 | 98.3 |
| 637 | 10 | 86.5 | 8.8 | 74.8 | 78.1 | 88.6 | 94.4 | 98.2 |
| 286104 | 15 | 92.9 | 2.2 | 90.4 | 91.4 | 92 | 94.4 | 98.3 |
| 154116 | 1 | 84.3 | nan | 84.3 | 84.3 | 84.3 | 84.3 | 84.3 |
| 1912215 | 3 | 93.1 | 3 | 91.1 | 91.3 | 91.6 | 94 | 96.5 |
| 157920 | 10 | 95.5 | 4.4 | 90.4 | 90.5 | 97.8 | 99.2 | 99.8 |
| 413496 | 1 | 99.3 | nan | 99.3 | 99.3 | 99.3 | 99.3 | 99.3 |
| 1906945 | 3 | 94.7 | 2.4 | 93.2 | 93.3 | 93.4 | 95.4 | 97.4 |
| 414371 | 10 | 86.2 | 2.1 | 83.7 | 84.8 | 86.2 | 87 | 90.8 |
| 2767842 | 6 | 97.5 | 1.2 | 96.6 | 96.7 | 97.2 | 97.8 | 99.7 |
| 2812025 | 1 | 81.8 | nan | 81.8 | 81.8 | 81.8 | 81.8 | 81.8 |
| 222 | 36 | 96.5 | 1.7 | 94 | 95 | 97.1 | 97.5 | 99.7 |
| 2767889 | 1 | 88 | nan | 88 | 88 | 88 | 88 | 88 |
| 572511 | 6 | 86.2 | 6 | 82 | 82.7 | 84.2 | 85.8 | 98 |
| 73029 | 1 | 93.7 | nan | 93.7 | 93.7 | 93.7 | 93.7 | 93.7 |
| 2129 | 1 | 95.3 | nan | 95.3 | 95.3 | 95.3 | 95.3 | 95.3 |
| 28453 | 91 | 91.4 | 3.4 | 88.6 | 89 | 89.8 | 93.8 | 99.8 |
| 29521 | 1 | 97.6 | nan | 97.6 | 97.6 | 97.6 | 97.6 | 97.6 |
| 43668 | 15 | 92 | 2.4 | 89.2 | 89.9 | 91.3 | 93.7 | 96.5 |
| 32257 | 3 | 88.5 | 1.7 | 87 | 87.5 | 88 | 89.2 | 90.4 |
| 261963 | 3 | 90.8 | 2.7 | 89.1 | 89.3 | 89.5 | 91.7 | 93.9 |
| 85413 | 21 | 92.3 | 1.7 | 90.4 | 91.2 | 91.6 | 93.6 | 95.9 |

| | | | | | | | | |
|---|---|---|---|---|---|---|---|---|
| 2782231 | 21 | 92.9 | 1.6 | 90.5 | 91.5 | 93 | 94.1 | 96 |
| 326319 | 3 | 97 | 1.9 | 95.8 | 95.8 | 95.9 | 97.6 | 99.2 |
| 1101 | 1 | 81.8 | nan | 81.8 | 81.8 | 81.8 | 81.8 | 81.8 |
| 1335745 | 1 | 83.8 | nan | 83.8 | 83.8 | 83.8 | 83.8 | 83.8 |
| 42255 | 6 | 80.5 | 4.2 | 77.5 | 78.7 | 79.2 | 79.6 | 88.9 |
| 1621534 | 3 | 89.7 | 5 | 86.7 | 86.8 | 87 | 91.2 | 95.5 |
| 568 | 3 | 99.3 | 0.6 | 98.9 | 99 | 99 | 99.5 | 99.9 |
| 83681 | 1 | 88.4 | nan | 88.4 | 88.4 | 88.4 | 88.4 | 88.4 |
| 1379908 | 3 | 93.5 | 1.3 | 92.5 | 92.8 | 93 | 94 | 95 |
| 204286 | 21 | 98.2 | 1.2 | 96.1 | 96.8 | 98.5 | 99.1 | 99.8 |
| 1754 | 1 | 95.7 | nan | 95.7 | 95.7 | 95.7 | 95.7 | 95.7 |
| 35798 | 1 | 90.5 | nan | 90.5 | 90.5 | 90.5 | 90.5 | 90.5 |
| 635 | 15 | 98.5 | 0.8 | 97.5 | 97.8 | 98.1 | 99.1 | 100 |
| 239934 | 1 | 80.7 | nan | 80.7 | 80.7 | 80.7 | 80.7 | 80.7 |
| 1854 | 3 | 84 | 1.4 | 83 | 83.2 | 83.4 | 84.5 | 85.6 |
| 28228 | 6 | 89.5 | 1.4 | 88 | 88.5 | 89.2 | 90.1 | 91.7 |
| 1522056 | 1 | 93 | nan | 93 | 93 | 93 | 93 | 93 |
| 1253 | 15 | 87.1 | 5.5 | 82.7 | 83.1 | 83.3 | 89.9 | 99.9 |
| 2767887 | 15 | 86.1 | 4.3 | 81 | 83.7 | 84.2 | 87.9 | 99.3 |
| 1647 | 6 | 80.1 | 6 | 74.4 | 74.8 | 79.6 | 84.4 | 87.9 |
| 1730 | 3 | 98.7 | 0.5 | 98.3 | 98.4 | 98.5 | 98.8 | 99.2 |
| 47420 | 28 | 91 | 2.4 | 87.4 | 89.6 | 90.2 | 91.3 | 97.4 |
| 13134 | 3 | 89.4 | 7.7 | 85 | 85 | 85 | 91.7 | 98.3 |
| 39948 | 3 | 86.9 | 11.2 | 80.4 | 80.4 | 80.4 | 90.1 | 99.8 |
| 2039639 | 3 | 91 | 0.7 | 90.3 | 90.7 | 91 | 91.3 | 91.7 |
| 1269 | 3 | 95.5 | 3.2 | 93.5 | 93.6 | 93.7 | 96.5 | 99.2 |
| 75309 | 1 | 94.2 | nan | 94.2 | 94.2 | 94.2 | 94.2 | 94.2 |

| | | | | | | | | |
|---|---|---|---|---|---|---|---|---|
| 2060 | 1 | 97.9 | nan | 97.9 | 97.9 | 97.9 | 97.9 | 97.9 |
| 191 | 28 | 95.2 | 3.2 | 91.3 | 91.7 | 96.9 | 98 | 99.3 |
| 2767881 | 3 | 85.3 | 3.5 | 82.5 | 83.3 | 84.2 | 86.7 | 89.2 |
| 83461 | 1 | 89.3 | nan | 89.3 | 89.3 | 89.3 | 89.3 | 89.3 |
| 146937 | 3 | 85.8 | 1.3 | 84.3 | 85.2 | 86.1 | 86.5 | 86.9 |
| 1705353 | 3 | 96.9 | 1.4 | 96 | 96.1 | 96.2 | 97.4 | 98.6 |
| 1163 | 1 | 97.4 | nan | 97.4 | 97.4 | 97.4 | 97.4 | 97.4 |
| 354349 | 3 | 99.8 | 0 | 99.8 | 99.8 | 99.8 | 99.8 | 99.8 |
| 225842 | 3 | 85.4 | 1.4 | 84.2 | 84.6 | 85 | 86 | 86.9 |
| 168934 | 3 | 92.1 | 1.8 | 90.7 | 91.1 | 91.5 | 92.8 | 94.2 |
| 2034 | 28 | 96.4 | 1.3 | 94.2 | 95.4 | 96.5 | 96.8 | 99.2 |
| 291183 | 6 | 92.3 | 2 | 90.4 | 90.6 | 91.9 | 94.2 | 94.6 |
| 626119 | 1 | 83.6 | nan | 83.6 | 83.6 | 83.6 | 83.6 | 83.6 |
| 2685905 | 1 | 94.6 | nan | 94.6 | 94.6 | 94.6 | 94.6 | 94.6 |
| 79328 | 10 | 87 | 3.5 | 84.5 | 85.2 | 86 | 87.6 | 96.3 |
| 182709 | 1 | 84.8 | nan | 84.8 | 84.8 | 84.8 | 84.8 | 84.8 |
| 110932 | 3 | 93.6 | 0.9 | 93 | 93.1 | 93.2 | 94 | 94.7 |
| 219181 | 1 | 91.3 | nan | 91.3 | 91.3 | 91.3 | 91.3 | 91.3 |
| 263377 | 3 | 91.5 | 1.5 | 90.6 | 90.7 | 90.7 | 92 | 93.2 |
| 1743 | 3 | 87.4 | 4.3 | 84.7 | 85 | 85.2 | 88.8 | 92.4 |
| 890 | 6 | 82.5 | 0.9 | 81.1 | 82.2 | 82.6 | 83.1 | 83.6 |
| 45667 | 3 | 90.4 | 0.9 | 89.7 | 89.9 | 90.1 | 90.8 | 91.5 |
| 168657 | 6 | 100 | 0 | 100 | 100 | 100 | 100 | 100 |
| 84565 | 1 | 86.1 | nan | 86.1 | 86.1 | 86.1 | 86.1 | 86.1 |
| 375288 | 6 | 95.5 | 2 | 94 | 94.5 | 94.8 | 95.2 | 99.4 |
| 34067 | 1 | 98.7 | nan | 98.7 | 98.7 | 98.7 | 98.7 | 98.7 |
| 265570 | 3 | 93.7 | 3.3 | 91.7 | 91.8 | 91.8 | 94.7 | 97.5 |

| | | | | | | | | |
|---|---|---|---|---|---|---|---|---|
| 56112 | 3 | 99.2 | 0.6 | 98.9 | 98.9 | 98.9 | 99.4 | 99.9 |
| 29487 | 3 | 97.7 | 0.2 | 97.5 | 97.6 | 97.7 | 97.8 | 97.8 |
| 970 | 3 | 95.4 | 3.3 | 93.4 | 93.5 | 93.5 | 96.3 | 99.2 |
| 51668 | 3 | 91.8 | 0.4 | 91.5 | 91.5 | 91.6 | 91.9 | 92.3 |
| 1434046 | 10 | 87.6 | 7.6 | 79 | 79.2 | 90.3 | 94.4 | 96.3 |
| 2370 | 1 | 92.2 | nan | 92.2 | 92.2 | 92.2 | 92.2 | 92.2 |
| 1330547 | 28 | 97.9 | 0.9 | 97 | 97.2 | 97.5 | 98.4 | 99.8 |
| 1742989 | 10 | 94.4 | 2.9 | 91.3 | 91.5 | 94.8 | 96.2 | 99.8 |
| 92944 | 1 | 93.9 | nan | 93.9 | 93.9 | 93.9 | 93.9 | 93.9 |
| 10 | 3 | 91.7 | 5.2 | 88.5 | 88.7 | 88.8 | 93.2 | 97.7 |
| 231454 | 6 | 89.7 | 1.7 | 87.6 | 88.4 | 89.9 | 90.4 | 92.2 |
| 198251 | 1 | 59.3 | nan | 59.3 | 59.3 | 59.3 | 59.3 | 59.3 |
| 293088 | 10 | 93.8 | 2.8 | 92.1 | 92.4 | 92.6 | 92.9 | 100 |
| 1742992 | 3 | 97.9 | 1.8 | 96.8 | 96.8 | 96.9 | 98.4 | 99.9 |
| 2321115 | 3 | 90 | 1.3 | 89.2 | 89.2 | 89.3 | 90.4 | 91.5 |
| 150247 | 21 | 94.5 | 2.8 | 92 | 92.1 | 94.2 | 95 | 100 |
| 1295327 | 1 | 89.2 | nan | 89.2 | 89.2 | 89.2 | 89.2 | 89.2 |
| 111500 | 3 | 92.6 | 4.5 | 89.8 | 90 | 90.2 | 94 | 97.7 |
| 28263 | 3 | 94.1 | 0.2 | 93.9 | 94 | 94 | 94.2 | 94.3 |
| 1470540 | 10 | 90 | 4 | 86.3 | 87.1 | 88.8 | 91.9 | 99.2 |
| 93 | 3 | 92.3 | 1.3 | 91.2 | 91.6 | 91.9 | 92.8 | 93.7 |
| 872 | 28 | 77.9 | 9.9 | 65.7 | 67.9 | 81.5 | 86.6 | 93.7 |
| 2071 | 1 | 92.3 | nan | 92.3 | 92.3 | 92.3 | 92.3 | 92.3 |
| 2800686 | 3 | 88.3 | 2.5 | 86.2 | 86.8 | 87.5 | 89.3 | 91.1 |
| 37914 | 10 | 91.1 | 5.8 | 84.3 | 84.5 | 94.2 | 94.4 | 98.7 |
| 2801844 | 1 | 87.5 | nan | 87.5 | 87.5 | 87.5 | 87.5 | 87.5 |
| 28884 | 3 | 88.4 | 2.9 | 86.5 | 86.7 | 86.9 | 89.3 | 91.7 |

| | | | | | | | | |
|---|---|---|---|---|---|---|---|---|
| 71655 | 3 | 96.1 | 2.4 | 94.7 | 94.7 | 94.7 | 96.8 | 98.8 |
| 1111 | 1 | 93.8 | nan | 93.8 | 93.8 | 93.8 | 93.8 | 93.8 |
| 1843210 | 1 | 82 | nan | 82 | 82 | 82 | 82 | 82 |
| 1778653 | 3 | 88.1 | 2.5 | 85.6 | 86.8 | 88.1 | 89.3 | 90.5 |
| 55079 | 3 | 92.7 | 6.3 | 89.1 | 89.1 | 89.1 | 94.5 | 100 |
| 2680004 | 3 | 94.1 | 0.7 | 93.3 | 93.8 | 94.2 | 94.5 | 94.7 |
| 162289 | 3 | 74.1 | 3 | 72.1 | 72.4 | 72.7 | 75.1 | 77.5 |
| 29330 | 3 | 76.3 | 1.2 | 75.1 | 75.7 | 76.2 | 76.8 | 77.5 |
| 475 | 21 | 85.7 | 5.8 | 77.3 | 78.5 | 86.8 | 89.1 | 98.3 |
| 105 | 1 | 92.8 | nan | 92.8 | 92.8 | 92.8 | 92.8 | 92.8 |
| 76634 | 3 | 87.7 | 3.6 | 85.3 | 85.6 | 85.9 | 88.8 | 91.8 |
| 323449 | 10 | 91.4 | 2 | 89 | 90.3 | 90.6 | 92.9 | 95.1 |
| 51366 | 1 | 99.9 | nan | 99.9 | 99.9 | 99.9 | 99.9 | 99.9 |
| 1505663 | 1 | 66 | nan | 66 | 66 | 66 | 66 | 66 |
| 1330546 | 1 | 95.8 | nan | 95.8 | 95.8 | 95.8 | 95.8 | 95.8 |
| 400947 | 3 | 97.2 | 1.9 | 96.1 | 96.1 | 96.1 | 97.8 | 99.4 |
| 2834348 | 1 | 89.7 | nan | 89.7 | 89.7 | 89.7 | 89.7 | 89.7 |
| 245876 | 3 | 88.1 | 6.7 | 84.2 | 84.2 | 84.2 | 90 | 95.8 |
| 1869285 | 3 | 94 | 2.9 | 92.3 | 92.3 | 92.3 | 94.8 | 97.3 |
| 511745 | 1 | 89.3 | nan | 89.3 | 89.3 | 89.3 | 89.3 | 89.3 |
| 49082 | 1 | 91 | nan | 91 | 91 | 91 | 91 | 91 |
| 265976 | 1 | 94.9 | nan | 94.9 | 94.9 | 94.9 | 94.9 | 94.9 |
| 106591 | 6 | 93.2 | 2.2 | 90 | 91.8 | 94 | 94.7 | 95.5 |
| 2675230 | 6 | 88.6 | 2.3 | 86.1 | 87.2 | 88.3 | 89.2 | 92.6 |
| 32 | 1 | 95.2 | nan | 95.2 | 95.2 | 95.2 | 95.2 | 95.2 |
| 1073531 | 3 | 94.9 | 1 | 94.3 | 94.3 | 94.3 | 95.2 | 96 |
| 2060094 | 1 | 93 | nan | 93 | 93 | 93 | 93 | 93 |

| | | | | | | | | |
|---|---|---|---|---|---|---|---|---|
| 1221880 | 3 | 92 | 2.1 | 90.6 | 90.8 | 91.1 | 92.8 | 94.4 |
| 1848399 | 3 | 91.1 | 1.7 | 89.6 | 90.2 | 90.7 | 91.8 | 93 |
| 1960084 | 1 | 94.4 | nan | 94.4 | 94.4 | 94.4 | 94.4 | 94.4 |
| 551759 | 3 | 87.3 | 3.8 | 85.1 | 85.2 | 85.2 | 88.5 | 91.7 |
| 2039723 | 3 | 87.2 | 1.2 | 86 | 86.6 | 87.2 | 87.8 | 88.4 |
| 305976 | 1 | 86.3 | nan | 86.3 | 86.3 | 86.3 | 86.3 | 86.3 |
| 57495 | 1 | 93.8 | nan | 93.8 | 93.8 | 93.8 | 93.8 | 93.8 |
| 73918 | 1 | 90.9 | nan | 90.9 | 90.9 | 90.9 | 90.9 | 90.9 |
| 46254 | 3 | 91.7 | 4.2 | 89.2 | 89.2 | 89.3 | 92.9 | 96.5 |
| 507 | 3 | 98.9 | 0.6 | 98.5 | 98.5 | 98.6 | 99.1 | 99.6 |

Table S5. PID in 747 species

| Species | count | mean | std | min | 25% | 50% | 75% | max |
|---|---|---|---|---|---|---|---|---|
| 103855 | 153 | 99.9 | 0 | 99.9 | 99.9 | 99.9 | 100 | 100 |
| 104628 | 10 | 99.8 | 0 | 99.8 | 99.8 | 99.8 | 99.9 | 99.9 |
| 1063 | 15 | 98.4 | 1.9 | 95.8 | 95.9 | 99.1 | 100 | 100 |
| 106590 | 3 | 99 | 0.4 | 98.7 | 98.8 | 98.8 | 99.1 | 99.4 |
| 106592 | 15 | 97 | 2.3 | 94.6 | 94.6 | 96.2 | 99 | 100 |
| 487 | 4186 | 99.4 | 0.3 | 98.9 | 99.2 | 99.3 | 99.8 | 100 |
| 1428 | 528 | 96.2 | 2.4 | 91.2 | 94.8 | 95.9 | 98.5 | 100 |
| 1903056 | 3 | 97.2 | 2.3 | 95.9 | 95.9 | 95.9 | 97.9 | 99.9 |
| 314275 | 36 | 99.6 | 0.1 | 99.4 | 99.5 | 99.6 | 99.7 | 100 |
| 1905730 | 78 | 99.9 | 0 | 99.9 | 99.9 | 99.9 | 99.9 | 100 |
| 38301 | 10 | 99.6 | 0.5 | 99.1 | 99.1 | 100 | 100 | 100 |
| 38313 | 153 | 99.5 | 0.3 | 99.1 | 99.3 | 99.6 | 99.8 | 99.9 |
| 669 | 15 | 99.1 | 1.2 | 97.4 | 97.4 | 99.9 | 100 | 100 |
| 669502 | 3 | 99.9 | 0.1 | 99.9 | 99.9 | 99.9 | 100 | 100 |
| 670 | 1431 | 99.8 | 0.1 | 99.5 | 99.8 | 99.9 | 99.9 | 100 |
| 518 | 136 | 99.7 | 0.3 | 99 | 99.6 | 99.8 | 99.9 | 100 |
| 569 | 10 | 99.9 | 0.1 | 99.8 | 99.8 | 99.8 | 100 | 100 |
| 571 | 351 | 99.6 | 0.3 | 98.8 | 99.5 | 99.6 | 99.9 | 100 |
| 80866 | 6 | 99.8 | 0.2 | 99.6 | 99.6 | 99.8 | 99.9 | 99.9 |
| 813 | 703 | 99.5 | 0.3 | 99.2 | 99.3 | 99.3 | 99.9 | 100 |
| 60552 | 15 | 99.9 | 0.1 | 99.8 | 99.8 | 99.9 | 100 | 100 |
| 60890 | 21 | 97.2 | 3.8 | 91.4 | 91.4 | 99.1 | 100 | 100 |
| 40041 | 210 | 99.3 | 0.1 | 99 | 99.3 | 99.4 | 99.4 | 100 |
| 1280 | 158766 | 99.8 | 0.3 | 95.5 | 99.7 | 99.8 | 99.9 | 100 |
| 1679 | 78 | 99.7 | 0.1 | 99.5 | 99.7 | 99.7 | 99.8 | 100 |

| | | | | | | | | |
|---|---|---|---|---|---|---|---|---|
| 64187 | 2278 | 99.4 | 0.6 | 98.7 | 98.8 | 99.9 | 100 | 100 |
| 35554 | 3 | 99.4 | 0.4 | 99.2 | 99.2 | 99.2 | 99.6 | 99.9 |
| 35703 | 21 | 99.9 | 0.1 | 99.7 | 99.7 | 99.9 | 100 | 100 |
| 730 | 55 | 99.7 | 0.3 | 99.3 | 99.3 | 99.9 | 100 | 100 |
| 731 | 351 | 99.7 | 0.2 | 99.2 | 99.5 | 99.6 | 100 | 100 |
| 380021 | 105 | 97.2 | 2.9 | 89.3 | 93.6 | 99.1 | 99.9 | 100 |
| 382 | 153 | 99.9 | 0.1 | 99.8 | 99.9 | 99.9 | 100 | 100 |
| 384 | 21 | 95.6 | 2.5 | 90.7 | 94.4 | 95 | 97.9 | 99.6 |
| 396 | 91 | 99.7 | 0.1 | 99.5 | 99.6 | 99.7 | 99.8 | 100 |
| 40214 | 55 | 99.5 | 0.1 | 99.2 | 99.4 | 99.5 | 99.6 | 99.7 |
| 40324 | 946 | 99 | 0.7 | 96.9 | 98.9 | 99.1 | 99.3 | 100 |
| 40477 | 190 | 99.8 | 0.2 | 98.9 | 99.8 | 99.9 | 99.9 | 100 |
| 40480 | 153 | 99.9 | 0 | 99.8 | 99.9 | 99.9 | 99.9 | 100 |
| 90105 | 45 | 100 | 0 | 99.9 | 99.9 | 100 | 100 | 100 |
| 519 | 3403 | 100 | 0.1 | 99.7 | 100 | 100 | 100 | 100 |
| 28035 | 153 | 99.9 | 0.1 | 99.8 | 99.9 | 99.9 | 100 | 100 |
| 28037 | 28 | 97.9 | 2.1 | 94.2 | 97.7 | 99.1 | 99.2 | 100 |
| 28038 | 55 | 99.6 | 0.1 | 99.3 | 99.6 | 99.6 | 99.7 | 99.8 |
| 87883 | 171 | 98.8 | 3.3 | 89.1 | 99.9 | 99.9 | 99.9 | 100 |
| 115981 | 10 | 99.9 | 0 | 99.9 | 99.9 | 99.9 | 99.9 | 100 |
| 1160769 | 10 | 100 | 0 | 100 | 100 | 100 | 100 | 100 |
| 550 | 253 | 99.2 | 0.3 | 98.8 | 98.9 | 99.2 | 99.4 | 100 |
| 539813 | 6 | 99.3 | 0.5 | 98.8 | 99 | 99.3 | 99.6 | 100 |
| 2259622 | 10 | 99.8 | 0.1 | 99.6 | 99.7 | 99.8 | 99.8 | 100 |
| 208962 | 210 | 99.9 | 0 | 99.8 | 99.9 | 99.9 | 99.9 | 100 |
| 386891 | 21 | 99 | 0.7 | 98.2 | 98.3 | 99.1 | 100 | 100 |
| 387 | 3 | 95.8 | 2.6 | 93.8 | 94.3 | 94.9 | 96.8 | 98.8 |

| | | | | | | | | |
|---|---|---|---|---|---|---|---|---|
| 291644 | 3 | 99.9 | 0 | 99.9 | 99.9 | 99.9 | 99.9 | 99.9 |
| 29459 | 1540 | 100 | 0 | 99.8 | 99.9 | 100 | 100 | 100 |
| 2098 | 91 | 99.7 | 0.1 | 99.6 | 99.7 | 99.7 | 99.8 | 99.8 |
| 2099 | 3 | 99.6 | 0.1 | 99.5 | 99.5 | 99.6 | 99.6 | 99.6 |
| 210 | 13041 | 98.4 | 0.5 | 95 | 98.1 | 98.3 | 98.6 | 100 |
| 1079 | 3 | 99.5 | 0.2 | 99.3 | 99.4 | 99.6 | 99.7 | 99.7 |
| 470 | 33670 | 99.8 | 0.5 | 93.1 | 99.8 | 99.8 | 99.9 | 100 |
| 633 | 91 | 99.9 | 0 | 99.9 | 99.9 | 99.9 | 99.9 | 100 |
| 103796 | 15 | 100 | 0 | 99.9 | 99.9 | 100 | 100 | 100 |
| 103816 | 15 | 99.9 | 0.1 | 99.8 | 99.8 | 99.9 | 99.9 | 99.9 |
| 1334 | 21 | 98.9 | 0.4 | 98.4 | 98.5 | 99 | 99.2 | 99.7 |
| 1579 | 15 | 99.9 | 0.1 | 99.6 | 99.8 | 99.8 | 100 | 100 |
| 446 | 1485 | 99.4 | 0.6 | 97.2 | 99.5 | 99.5 | 99.9 | 100 |
| 696485 | 6 | 98.2 | 1.3 | 97.3 | 97.3 | 97.4 | 99.3 | 100 |
| 1680 | 28 | 99.5 | 0.3 | 99 | 99.5 | 99.6 | 99.7 | 99.8 |
| 1681 | 21 | 99.6 | 0.4 | 99 | 99.2 | 99.8 | 99.8 | 100 |
| 1767 | 91 | 99.8 | 0.1 | 99.6 | 99.7 | 99.8 | 99.9 | 100 |
| 1770 | 45 | 99.9 | 0.1 | 99.7 | 99.9 | 99.9 | 100 | 100 |
| 190893 | 3 | 99.8 | 0.1 | 99.8 | 99.8 | 99.8 | 99.8 | 99.9 |
| 192 | 21 | 99.4 | 0.4 | 98.9 | 99.1 | 99.2 | 99.8 | 100 |
| 1922217 | 36 | 79.9 | 7.9 | 72.6 | 73.2 | 77.2 | 87.8 | 100 |
| 192954 | 3 | 100 | 0 | 100 | 100 | 100 | 100 | 100 |
| 195 | 903 | 99.5 | 0.3 | 98.3 | 99.3 | 99.6 | 99.8 | 100 |
| 138074 | 3 | 91.6 | 7.2 | 87.4 | 87.4 | 87.4 | 93.7 | 99.9 |
| 40215 | 28 | 99.7 | 0.1 | 99.5 | 99.6 | 99.7 | 99.7 | 99.9 |
| 576611 | 10 | 95.6 | 3.1 | 92.6 | 92.9 | 94.9 | 98.6 | 100 |
| 57706 | 15 | 99.9 | 0.1 | 99.6 | 99.7 | 99.9 | 100 | 100 |

| | | | | | | | | |
|---|---|---|---|---|---|---|---|---|
| 573 | 314028 | 99.7 | 0.5 | 97.4 | 99.9 | 99.9 | 99.9 | 100 |
| 1117645 | 171 | 99.8 | 0.2 | 99 | 99.8 | 99.8 | 99.9 | 100 |
| 1311 | 5253 | 99.6 | 0.3 | 98.3 | 99.5 | 99.5 | 99.7 | 100 |
| 1402 | 465 | 100 | 0.1 | 99.5 | 100 | 100 | 100 | 100 |
| 158836 | 5151 | 99.7 | 0.5 | 96.2 | 99.7 | 99.8 | 99.9 | 100 |
| 1613 | 378 | 99.6 | 0.4 | 98 | 99.5 | 99.7 | 99.8 | 100 |
| 1648 | 15 | 99.9 | 0.2 | 99.6 | 99.7 | 99.9 | 100 | 100 |
| 1719 | 4656 | 99.9 | 0.1 | 99.4 | 99.8 | 99.9 | 100 | 100 |
| 1778540 | 10 | 99.8 | 0.2 | 99.6 | 99.6 | 99.8 | 100 | 100 |
| 2058152 | 91 | 99.9 | 0 | 99.9 | 99.9 | 99.9 | 100 | 100 |
| 243161 | 66 | 100 | 0 | 100 | 100 | 100 | 100 | 100 |
| 264203 | 3 | 100 | 0 | 100 | 100 | 100 | 100 | 100 |
| 246432 | 28 | 99.8 | 0.1 | 99.6 | 99.8 | 99.8 | 99.9 | 100 |
| 246787 | 3 | 99.5 | 0.3 | 99.3 | 99.3 | 99.3 | 99.6 | 99.9 |
| 213 | 15 | 99.8 | 0.1 | 99.7 | 99.7 | 99.7 | 99.7 | 100 |
| 2133 | 6 | 100 | 0 | 100 | 100 | 100 | 100 | 100 |
| 1185650 | 190 | 100 | 0.1 | 99.8 | 99.9 | 100 | 100 | 100 |
| 650 | 6 | 99.8 | 0.1 | 99.7 | 99.8 | 99.8 | 99.9 | 99.9 |
| 65058 | 91 | 99.8 | 0.1 | 99.6 | 99.8 | 99.9 | 99.9 | 100 |
| 1772 | 6 | 99.8 | 0.1 | 99.7 | 99.7 | 99.7 | 99.9 | 100 |
| 1773 | 27495 | 99.9 | 0.2 | 97.7 | 99.9 | 99.9 | 100 | 100 |
| 2756 | 10 | 100 | 0.1 | 99.9 | 99.9 | 100 | 100 | 100 |
| 582 | 300 | 98.8 | 0.9 | 97 | 98 | 99.3 | 99.8 | 100 |
| 584 | 2346 | 99.9 | 0.1 | 99.7 | 99.9 | 99.9 | 99.9 | 100 |
| 1335 | 21 | 97.7 | 3.5 | 92.2 | 92.3 | 99.5 | 100 | 100 |
| 1492 | 66 | 98.3 | 1.9 | 95.9 | 96 | 99.8 | 100 | 100 |
| 149385 | 3 | 100 | 0 | 100 | 100 | 100 | 100 | 100 |

| | | | | | | | | |
|---|---|---|---|---|---|---|---|---|
| 149387 | 3 | 100 | 0 | 100 | 100 | 100 | 100 | 100 |
| 1493872 | 3 | 99.9 | 0.1 | 99.8 | 99.8 | 99.8 | 99.9 | 100 |
| 149390 | 3 | 100 | 0 | 100 | 100 | 100 | 100 | 100 |
| 1530123 | 190 | 99.8 | 0.2 | 99.4 | 99.7 | 99.8 | 100 | 100 |
| 334542 | 15 | 99.9 | 0 | 99.9 | 99.9 | 99.9 | 99.9 | 100 |
| 258 | 3 | 99.2 | 0.2 | 99 | 99.1 | 99.2 | 99.2 | 99.3 |
| 139 | 21 | 99.6 | 0.2 | 99.4 | 99.4 | 99.5 | 99.5 | 100 |
| 1390 | 703 | 99.6 | 0.3 | 98.4 | 99.3 | 99.7 | 99.8 | 100 |
| 1238 | 2016 | 99.2 | 0.6 | 98.1 | 98.6 | 98.9 | 99.8 | 100 |
| 119857 | 91 | 99.2 | 1 | 97.7 | 98 | 99.9 | 99.9 | 100 |
| 120045 | 3 | 99.9 | 0.1 | 99.9 | 99.9 | 99.9 | 100 | 100 |
| 13373 | 231 | 100 | 0 | 100 | 100 | 100 | 100 | 100 |
| 1340 | 6 | 99.5 | 0.4 | 99 | 99.1 | 99.4 | 99.8 | 100 |
| 1351 | 1485 | 99.4 | 0.4 | 98.9 | 99 | 99.1 | 99.8 | 100 |
| 1352 | 24090 | 99.9 | 0.1 | 99.5 | 99.9 | 100 | 100 | 100 |
| 1354 | 66 | 99.9 | 0.1 | 99.8 | 99.8 | 99.9 | 99.9 | 100 |
| 135461 | 378 | 99.8 | 0.3 | 98.8 | 99.8 | 99.9 | 99.9 | 100 |
| 1355477 | 91 | 98.2 | 1.5 | 95.4 | 96.9 | 97.7 | 99.7 | 100 |
| 1359 | 91 | 99.8 | 0.2 | 99.2 | 99.7 | 99.8 | 99.9 | 100 |
| 1360 | 105 | 99.7 | 0.2 | 99.4 | 99.5 | 99.6 | 99.8 | 100 |
| 948564 | 3 | 100 | 0 | 100 | 100 | 100 | 100 | 100 |
| 95486 | 153 | 99 | 1.4 | 95.3 | 99.3 | 99.5 | 99.8 | 100 |
| 1295 | 21 | 98.5 | 2.1 | 95.2 | 95.3 | 99.7 | 100 | 100 |
| 1296 | 36 | 99.7 | 0.1 | 99.5 | 99.6 | 99.6 | 99.7 | 99.9 |
| 1309 | 120 | 99.5 | 0.2 | 99.2 | 99.4 | 99.5 | 99.6 | 100 |
| 1310 | 15 | 99.7 | 0.2 | 99.4 | 99.5 | 99.7 | 99.8 | 100 |
| 1781 | 6 | 99.2 | 0.3 | 98.8 | 99.1 | 99.2 | 99.4 | 99.8 |

| | | | | | | | | |
|---|---|---|---|---|---|---|---|---|
| 1452 | 6 | 99.9 | 0.1 | 99.8 | 99.8 | 99.9 | 99.9 | 99.9 |
| 145458 | 6 | 99.9 | 0.1 | 99.8 | 99.8 | 99.8 | 99.9 | 100 |
| 1134687 | 231 | 99.2 | 1.6 | 93 | 99.5 | 99.9 | 99.9 | 100 |
| 1717 | 28 | 99.6 | 0.3 | 99 | 99.5 | 99.7 | 99.7 | 99.8 |
| 1718 | 300 | 99.6 | 0.2 | 98.7 | 99.5 | 99.6 | 99.7 | 100 |
| 1642299 | 3 | 97.5 | 2.1 | 96.3 | 96.3 | 96.3 | 98.2 | 100 |
| 164514 | 36 | 98.5 | 0.9 | 97.3 | 97.7 | 97.8 | 99.4 | 99.9 |
| 1705 | 3 | 99.8 | 0.2 | 99.7 | 99.8 | 99.8 | 99.9 | 100 |
| 299583 | 171 | 100 | 0.1 | 99.6 | 100 | 100 | 100 | 100 |
| 299766 | 21 | 99.9 | 0 | 99.9 | 99.9 | 99.9 | 99.9 | 100 |
| 2733571 | 6 | 99.5 | 0.4 | 99 | 99.1 | 99.5 | 99.8 | 99.9 |
| 1648923 | 66 | 99.7 | 0.3 | 98.9 | 99.6 | 99.9 | 99.9 | 100 |
| 1396 | 1596 | 96 | 2.5 | 91.4 | 94.4 | 95.9 | 98.4 | 100 |
| 1869227 | 3 | 70.1 | 25.9 | 55.2 | 55.2 | 55.2 | 77.6 | 100 |
| 28116 | 10 | 99.8 | 0.1 | 99.6 | 99.7 | 99.7 | 99.8 | 99.9 |
| 28129 | 6 | 99.6 | 0.1 | 99.6 | 99.6 | 99.7 | 99.7 | 99.7 |
| 28131 | 55 | 99.4 | 0.2 | 99.1 | 99.3 | 99.4 | 99.5 | 100 |
| 201 | 3 | 97.7 | 1.1 | 97 | 97 | 97.1 | 98 | 99 |
| 204039 | 66 | 99.9 | 0.2 | 99.6 | 100 | 100 | 100 | 100 |
| 180434 | 66 | 99.2 | 0.9 | 97.1 | 98.1 | 99.8 | 99.9 | 100 |
| 1806 | 10 | 99.8 | 0.3 | 99.4 | 99.4 | 100 | 100 | 100 |
| 204042 | 21 | 98.3 | 1 | 97.1 | 97.5 | 97.9 | 99.2 | 99.9 |
| 511 | 45 | 99.2 | 0.5 | 98.2 | 98.9 | 99.1 | 99.7 | 100 |
| 511145 | 28 | 100 | 0 | 100 | 100 | 100 | 100 | 100 |
| 2026188 | 10 | 95.3 | 3.6 | 92.2 | 92.2 | 93 | 99.1 | 99.8 |
| 1884905 | 6 | 99.3 | 0.1 | 99.2 | 99.3 | 99.4 | 99.4 | 99.4 |
| 192955 | 10 | 100 | 0 | 100 | 100 | 100 | 100 | 100 |

| | | | | | | | | |
|---|---|---|---|---|---|---|---|---|
| 196024 | 6 | 99.8 | 0.1 | 99.7 | 99.7 | 99.8 | 99.8 | 99.8 |
| 1962118 | 10 | 99.8 | 0.2 | 99.7 | 99.7 | 99.7 | 100 | 100 |
| 44688 | 6 | 99.9 | 0.1 | 99.9 | 99.9 | 100 | 100 | 100 |
| 450 | 15 | 99.9 | 0.1 | 99.8 | 99.9 | 99.9 | 100 | 100 |
| 134821 | 6 | 99.8 | 0.1 | 99.7 | 99.7 | 99.8 | 99.8 | 100 |
| 1653831 | 190 | 75.3 | 6.8 | 68 | 69.8 | 73.3 | 78.3 | 97.8 |
| 1685 | 630 | 99.6 | 0.4 | 97.8 | 99.6 | 99.7 | 99.8 | 100 |
| 169679 | 6 | 100 | 0 | 100 | 100 | 100 | 100 | 100 |
| 529 | 6 | 99.7 | 0.2 | 99.5 | 99.5 | 99.6 | 99.8 | 100 |
| 235 | 66 | 99.9 | 0 | 99.9 | 99.9 | 99.9 | 100 | 100 |
| 930166 | 6 | 99.4 | 0.3 | 99 | 99.2 | 99.4 | 99.5 | 100 |
| 217203 | 6 | 98.4 | 1.3 | 97.1 | 97.2 | 98.2 | 99.5 | 99.9 |
| 217204 | 3 | 98.1 | 1.4 | 97.2 | 97.3 | 97.4 | 98.6 | 99.8 |
| 2151 | 21 | 99.3 | 0.4 | 98.9 | 99 | 99.1 | 99.7 | 99.8 |
| 216816 | 171 | 99.5 | 0.4 | 98.4 | 99.4 | 99.7 | 99.8 | 100 |
| 221822 | 190 | 99.3 | 0.3 | 98.6 | 99.1 | 99.3 | 99.4 | 100 |
| 222805 | 55 | 99.7 | 0.2 | 99.4 | 99.5 | 99.8 | 99.9 | 100 |
| 2231116 | 3 | 60.7 | 2.4 | 58.7 | 59.4 | 60 | 61.6 | 63.3 |
| 686 | 15 | 100 | 0 | 99.9 | 99.9 | 100 | 100 | 100 |
| 2527775 | 21 | 99.8 | 0.1 | 99.7 | 99.8 | 99.8 | 99.8 | 100 |
| 853 | 55 | 97.5 | 1.2 | 96.1 | 96.7 | 97.2 | 97.6 | 100 |
| 2663009 | 15 | 99.7 | 0.1 | 99.5 | 99.5 | 99.8 | 99.8 | 99.9 |
| 1254 | 253 | 99.7 | 0.2 | 99 | 99.7 | 99.8 | 99.8 | 100 |
| 1255 | 153 | 99.5 | 0.9 | 96.8 | 99.8 | 99.8 | 99.9 | 100 |
| 361101 | 3 | 99.4 | 0.2 | 99.2 | 99.3 | 99.5 | 99.5 | 99.6 |
| 363952 | 10 | 99.4 | 0.5 | 98.7 | 99 | 99.6 | 99.8 | 100 |
| 504 | 3 | 99.3 | 0.3 | 98.9 | 99.2 | 99.4 | 99.5 | 99.5 |

| | | | | | | | | |
|---|---|---|---|---|---|---|---|---|
| 2488639 | 15 | 99.9 | 0 | 99.8 | 99.9 | 99.9 | 99.9 | 99.9 |
| 1620419 | 3 | 100 | 0 | 100 | 100 | 100 | 100 | 100 |
| 1622 | 3 | 99.8 | 0.1 | 99.7 | 99.7 | 99.7 | 99.8 | 99.9 |
| 1624 | 21 | 99.7 | 0.1 | 99.5 | 99.6 | 99.7 | 99.8 | 99.8 |
| 162426 | 6 | 94.7 | 5.7 | 89.5 | 89.5 | 94.7 | 99.8 | 99.9 |
| 1744 | 210 | 99.8 | 0.1 | 99.4 | 99.7 | 99.8 | 99.9 | 100 |
| 39791 | 6 | 99.7 | 0.1 | 99.7 | 99.7 | 99.7 | 99.8 | 99.8 |
| 2702 | 3 | 96.8 | 2.6 | 95.3 | 95.3 | 95.4 | 97.6 | 99.8 |
| 1296536 | 28 | 99.9 | 0 | 99.9 | 99.9 | 99.9 | 100 | 100 |
| 260554 | 6 | 99.9 | 0 | 99.9 | 99.9 | 99.9 | 99.9 | 99.9 |
| 263 | 66 | 98.9 | 1.1 | 97.5 | 97.5 | 99.5 | 100 | 100 |
| 155322 | 15 | 96.9 | 2.2 | 93.1 | 96 | 96 | 98.5 | 100 |
| 1547 | 3 | 100 | 0 | 100 | 100 | 100 | 100 | 100 |
| 480 | 55 | 99.7 | 0.1 | 99.5 | 99.6 | 99.6 | 99.7 | 100 |
| 481146 | 3 | 99.7 | 0.1 | 99.7 | 99.7 | 99.7 | 99.8 | 99.8 |
| 482957 | 3 | 99.4 | 0.1 | 99.3 | 99.3 | 99.4 | 99.4 | 99.4 |
| 485 | 1176 | 99.8 | 0.2 | 99.3 | 99.8 | 99.9 | 99.9 | 100 |
| 28141 | 66 | 99.9 | 0 | 99.9 | 99.9 | 99.9 | 99.9 | 100 |
| 28150 | 120 | 99.9 | 0.1 | 99.9 | 99.9 | 99.9 | 100 | 100 |
| 283734 | 325 | 98.4 | 2 | 95.8 | 95.9 | 99.9 | 99.9 | 100 |
| 28450 | 3828 | 97.8 | 2.6 | 94.4 | 94.6 | 99.9 | 99.9 | 100 |
| 663 | 325 | 99.3 | 1.7 | 93.4 | 99.6 | 99.9 | 99.9 | 100 |
| 75105 | 6 | 100 | 0.1 | 99.9 | 99.9 | 100 | 100 | 100 |
| 414 | 3 | 99.8 | 0.2 | 99.7 | 99.7 | 99.7 | 99.8 | 100 |
| 192812 | 6 | 97.2 | 2.6 | 94.8 | 94.8 | 97.2 | 99.6 | 99.7 |
| 76758 | 10 | 99.6 | 0.3 | 99.3 | 99.3 | 99.8 | 99.8 | 100 |
| 76759 | 21 | 97.9 | 2.1 | 93.1 | 96 | 97 | 100 | 100 |

| | | | | | | | | |
|---|---|---|---|---|---|---|---|---|
| 70255 | 10 | 100 | 0 | 99.9 | 100 | 100 | 100 | 100 |
| 70775 | 3 | 96.8 | 1.3 | 95.3 | 96.3 | 97.4 | 97.6 | 97.8 |
| 72407 | 3081 | 99.7 | 0.5 | 98.2 | 99.9 | 99.9 | 99.9 | 100 |
| 727 | 1953 | 99 | 0.5 | 96.1 | 98.8 | 99.2 | 99.3 | 100 |
| 729 | 91 | 99.3 | 0.3 | 98.5 | 99.1 | 99.4 | 99.5 | 99.7 |
| 738 | 153 | 99.4 | 0.2 | 99 | 99.2 | 99.3 | 99.6 | 99.9 |
| 747 | 1596 | 99.6 | 0.4 | 98.6 | 99.7 | 99.7 | 99.9 | 100 |
| 76760 | 3 | 99.9 | 0 | 99.9 | 99.9 | 99.9 | 99.9 | 99.9 |
| 76802 | 6 | 99.9 | 0.1 | 99.9 | 99.9 | 99.9 | 100 | 100 |
| 621 | 91 | 99.8 | 0.1 | 99.5 | 99.8 | 99.8 | 99.9 | 100 |
| 622 | 120 | 99.7 | 0.2 | 99.5 | 99.6 | 99.6 | 99.9 | 100 |
| 561879 | 21 | 99.6 | 0.1 | 99.5 | 99.5 | 99.6 | 99.7 | 100 |
| 562 | 915981 | 99.9 | 0.2 | 96.5 | 99.9 | 99.9 | 99.9 | 100 |
| 1513890 | 66 | 98.2 | 1.7 | 96.2 | 96.5 | 99.6 | 99.9 | 100 |
| 1619313 | 3 | 99.9 | 0 | 99.9 | 99.9 | 99.9 | 99.9 | 99.9 |
| 162 | 21 | 99.9 | 0 | 99.9 | 99.9 | 99.9 | 99.9 | 100 |
| 47715 | 253 | 99.8 | 0.1 | 99.4 | 99.7 | 99.8 | 99.8 | 100 |
| 189426 | 3 | 97.8 | 1.6 | 96.8 | 96.9 | 97 | 98.3 | 99.7 |
| 253 | 36 | 98.8 | 1.3 | 97.1 | 97.2 | 99.9 | 99.9 | 100 |
| 1286 | 28 | 99.9 | 0 | 99.9 | 99.9 | 99.9 | 100 | 100 |
| 317013 | 55 | 100 | 0.1 | 99.8 | 100 | 100 | 100 | 100 |
| 147645 | 6 | 99.6 | 0.1 | 99.5 | 99.6 | 99.7 | 99.7 | 99.7 |
| 147802 | 15 | 99.8 | 0 | 99.8 | 99.8 | 99.8 | 99.8 | 99.9 |
| 375 | 3 | 99.3 | 0.6 | 98.9 | 98.9 | 98.9 | 99.5 | 100 |
| 375175 | 10 | 99.9 | 0.1 | 99.9 | 99.9 | 99.9 | 100 | 100 |
| 77038 | 3 | 93.2 | 5.7 | 89.9 | 89.9 | 89.9 | 94.8 | 99.8 |
| 777 | 15 | 99.9 | 0.1 | 99.8 | 99.8 | 99.9 | 100 | 100 |

| | | | | | | | | |
|---|---|---|---|---|---|---|---|---|
| 779 | 15 | 99.6 | 0.2 | 99.4 | 99.5 | 99.6 | 99.7 | 100 |
| 28080 | 3 | 99.1 | 0.3 | 98.9 | 99 | 99 | 99.2 | 99.4 |
| 28087 | 3 | 95.9 | 3.1 | 94 | 94.1 | 94.2 | 96.8 | 99.4 |
| 28090 | 21 | 99.6 | 0.1 | 99.4 | 99.5 | 99.5 | 99.6 | 99.8 |
| 28095 | 6 | 98 | 0.8 | 97.3 | 97.7 | 97.8 | 97.9 | 99.7 |
| 28108 | 3 | 99.2 | 0.5 | 98.8 | 98.9 | 99 | 99.3 | 99.7 |
| 1247 | 6 | 98.8 | 1.2 | 97.7 | 97.8 | 98.8 | 99.8 | 100 |
| 1689 | 3 | 99.8 | 0 | 99.8 | 99.8 | 99.8 | 99.8 | 99.8 |
| 1697053 | 3 | 99.6 | 0.3 | 99.4 | 99.4 | 99.4 | 99.7 | 99.9 |
| 197 | 13861 | 99.3 | 0.3 | 98.8 | 99.1 | 99.3 | 99.5 | 100 |
| 1245 | 66 | 99.6 | 0.5 | 98.4 | 99.7 | 99.8 | 99.9 | 100 |
| 1263550 | 28 | 100 | 0.1 | 99.9 | 99.9 | 100 | 100 | 100 |
| 119602 | 105 | 99.5 | 0.2 | 99.1 | 99.4 | 99.6 | 99.7 | 100 |
| 33967 | 10 | 99.2 | 0.6 | 98.4 | 98.6 | 99.6 | 99.7 | 99.8 |
| 33968 | 3 | 99.3 | 0.6 | 99 | 99 | 99 | 99.5 | 100 |
| 198620 | 3 | 99.2 | 0.2 | 99.1 | 99.1 | 99.1 | 99.3 | 99.5 |
| 199 | 105 | 98.2 | 0.7 | 96.4 | 98 | 98.1 | 98.7 | 100 |
| 109328 | 3 | 99.1 | 0.3 | 98.9 | 99 | 99 | 99.2 | 99.4 |
| 31963 | 3 | 100 | 0 | 100 | 100 | 100 | 100 | 100 |
| 319939 | 3 | 99.9 | 0.1 | 99.8 | 99.8 | 99.9 | 99.9 | 99.9 |
| 32002 | 10 | 98.9 | 1.3 | 97.3 | 97.4 | 99.8 | 99.9 | 100 |
| 33013 | 10 | 99.8 | 0.1 | 99.7 | 99.7 | 99.8 | 99.8 | 100 |
| 33014 | 3 | 99.9 | 0.1 | 99.8 | 99.8 | 99.9 | 99.9 | 99.9 |
| 33028 | 3 | 99.1 | 0.8 | 98.7 | 98.7 | 98.7 | 99.3 | 100 |
| 28144 | 3 | 99.9 | 0.1 | 99.9 | 99.9 | 99.9 | 100 | 100 |
| 33050 | 3 | 99.1 | 0.7 | 98.7 | 98.8 | 98.8 | 99.3 | 99.9 |
| 51663 | 10 | 99.4 | 0.3 | 99 | 99.1 | 99.4 | 99.5 | 100 |

| | | | | | | | | |
|---|---|---|---|---|---|---|---|---|
| 51665 | 3 | 99.4 | 0.3 | 99.2 | 99.2 | 99.2 | 99.5 | 99.8 |
| 51669 | 3 | 99.1 | 0.4 | 98.8 | 98.8 | 98.9 | 99.2 | 99.6 |
| 473423 | 78 | 99.8 | 0.2 | 99.6 | 99.6 | 99.8 | 100 | 100 |
| 47770 | 78 | 99.6 | 0.2 | 99.3 | 99.5 | 99.6 | 99.7 | 100 |
| 48296 | 325 | 98 | 1.9 | 95.7 | 95.9 | 99.6 | 99.7 | 100 |
| 34038 | 10 | 100 | 0.1 | 99.9 | 99.9 | 100 | 100 | 100 |
| 34062 | 45 | 98.9 | 0.4 | 98.3 | 98.5 | 98.9 | 99.2 | 100 |
| 1045010 | 28 | 100 | 0 | 100 | 100 | 100 | 100 | 100 |
| 676 | 55 | 99.8 | 0.1 | 99.8 | 99.8 | 99.8 | 99.9 | 100 |
| 67824 | 3 | 99.9 | 0.1 | 99.8 | 99.8 | 99.8 | 99.9 | 100 |
| 67351 | 10 | 100 | 0.1 | 99.9 | 99.9 | 100 | 100 | 100 |
| 674 | 21 | 99.7 | 0.2 | 99.4 | 99.5 | 99.8 | 99.8 | 99.9 |
| 675 | 3 | 99.9 | 0 | 99.9 | 99.9 | 99.9 | 99.9 | 99.9 |
| 1884913 | 3 | 99.3 | 0.2 | 99.1 | 99.2 | 99.3 | 99.3 | 99.4 |
| 1884914 | 3 | 99.5 | 0.1 | 99.4 | 99.5 | 99.6 | 99.6 | 99.6 |
| 1886 | 6 | 99.2 | 0.8 | 98.3 | 98.5 | 99.2 | 99.8 | 100 |
| 1270 | 36 | 98.6 | 1.3 | 96.1 | 98.7 | 99.2 | 99.3 | 100 |
| 29575 | 10 | 99.7 | 0.1 | 99.6 | 99.7 | 99.7 | 99.8 | 99.9 |
| 296 | 3 | 99.9 | 0 | 99.9 | 99.9 | 99.9 | 99.9 | 99.9 |
| 47763 | 6 | 95.2 | 1.7 | 93.3 | 94.1 | 94.9 | 96.1 | 97.9 |
| 43770 | 15 | 99 | 0.4 | 98.1 | 98.8 | 98.9 | 99.3 | 99.6 |
| 43771 | 3 | 99.8 | 0.1 | 99.7 | 99.8 | 99.8 | 99.8 | 99.9 |
| 1519 | 3 | 100 | 0 | 100 | 100 | 100 | 100 | 100 |
| 1520 | 15 | 95.4 | 3.7 | 89.2 | 91.6 | 96.7 | 98.1 | 99.6 |
| 1525 | 10 | 99.6 | 0.2 | 99.3 | 99.4 | 99.5 | 99.8 | 100 |
| 152500 | 3 | 100 | 0 | 100 | 100 | 100 | 100 | 100 |
| 1583 | 6 | 99.8 | 0.1 | 99.7 | 99.7 | 99.8 | 99.9 | 99.9 |

| | | | | | | | | |
|---|---|---|---|---|---|---|---|---|
| 1585 | 28 | 99.7 | 0.3 | 99.1 | 99.6 | 99.8 | 99.9 | 100 |
| 155892 | 6 | 98.9 | 0.8 | 98.2 | 98.2 | 98.8 | 99.5 | 100 |
| 1587 | 136 | 99.6 | 0.2 | 99.3 | 99.4 | 99.6 | 99.7 | 100 |
| 1829 | 3 | 99.5 | 0.4 | 99.3 | 99.3 | 99.3 | 99.7 | 100 |
| 1830 | 15 | 99.7 | 0.3 | 99.3 | 99.3 | 99.9 | 100 | 100 |
| 1833 | 3 | 99.8 | 0.1 | 99.8 | 99.8 | 99.8 | 99.8 | 99.9 |
| 1661 | 36 | 99.7 | 0.1 | 99.6 | 99.7 | 99.7 | 99.8 | 100 |
| 86040 | 15 | 99.9 | 0.1 | 99.8 | 99.8 | 99.9 | 100 | 100 |
| 86185 | 6 | 99.9 | 0.1 | 99.9 | 99.9 | 100 | 100 | 100 |
| 86192 | 105 | 98.4 | 1.6 | 96.1 | 96.5 | 99.3 | 99.9 | 100 |
| 82996 | 28 | 99.7 | 0.3 | 99.3 | 99.7 | 99.9 | 99.9 | 100 |
| 83655 | 66 | 99.4 | 0.8 | 97.1 | 99.1 | 99.9 | 99.9 | 100 |
| 83656 | 3 | 93 | 6.1 | 89.5 | 89.5 | 89.5 | 94.8 | 100 |
| 83684 | 3 | 99.7 | 0.1 | 99.6 | 99.7 | 99.7 | 99.7 | 99.7 |
| 837 | 136 | 99.7 | 0.1 | 99.6 | 99.7 | 99.7 | 99.8 | 100 |
| 299767 | 45 | 100 | 0 | 99.9 | 99.9 | 100 | 100 | 100 |
| 300 | 6 | 97.7 | 1.1 | 96.6 | 96.8 | 97.5 | 98.2 | 99.4 |
| 300181 | 10 | 99.9 | 0 | 99.9 | 99.9 | 99.9 | 99.9 | 99.9 |
| 303 | 351 | 97.6 | 1.7 | 90.7 | 97.2 | 97.7 | 98.6 | 100 |
| 519450 | 10 | 99 | 0.8 | 98 | 98.1 | 99.4 | 99.5 | 100 |
| 520 | 150426 | 100 | 0 | 99.3 | 100 | 100 | 100 | 100 |
| 243274 | 6 | 100 | 0 | 100 | 100 | 100 | 100 | 100 |
| 244320 | 15 | 99.4 | 0.8 | 98.3 | 98.3 | 100 | 100 | 100 |
| 244366 | 1081 | 99.8 | 0.2 | 99.3 | 99.6 | 99.9 | 99.9 | 100 |
| 1710 | 3 | 98.5 | 1.1 | 97.8 | 97.8 | 97.9 | 98.8 | 99.7 |
| 78327 | 3 | 99.9 | 0.1 | 99.8 | 99.8 | 99.8 | 99.9 | 100 |
| 784 | 15 | 98.7 | 0.3 | 98.1 | 98.3 | 98.9 | 98.9 | 99.2 |

| | | | | | | | | |
|---|---|---|---|---|---|---|---|---|
| 1148 | 10 | 100 | 0 | 100 | 100 | 100 | 100 | 100 |
| 439334 | 66 | 99.7 | 0.1 | 99.4 | 99.7 | 99.7 | 99.8 | 100 |
| 1915 | 3 | 99.5 | 0.3 | 99.3 | 99.3 | 99.3 | 99.5 | 99.8 |
| 45972 | 6 | 99.9 | 0.1 | 99.8 | 99.8 | 99.9 | 99.9 | 99.9 |
| 451 | 3 | 100 | 0.1 | 99.9 | 100 | 100 | 100 | 100 |
| 1303 | 45 | 98.7 | 0.6 | 97.2 | 98.1 | 99 | 99.3 | 99.4 |
| 2055 | 3 | 98.9 | 1 | 98.3 | 98.3 | 98.3 | 99.2 | 100 |
| 2055160 | 10 | 99.8 | 0.1 | 99.6 | 99.7 | 99.8 | 99.8 | 99.9 |
| 1173427 | 3 | 100 | 0 | 100 | 100 | 100 | 100 | 100 |
| 1736 | 3 | 98.8 | 0.9 | 98.3 | 98.3 | 98.3 | 99 | 99.8 |
| 651 | 36 | 99.7 | 0.1 | 99.5 | 99.6 | 99.7 | 99.7 | 100 |
| 37931 | 3 | 99.8 | 0.1 | 99.8 | 99.8 | 99.8 | 99.8 | 99.9 |
| 37734 | 15 | 99.8 | 0.1 | 99.8 | 99.8 | 99.8 | 99.9 | 100 |
| 70346 | 3 | 99.5 | 0.2 | 99.4 | 99.5 | 99.5 | 99.6 | 99.7 |
| 35790 | 78 | 100 | 0 | 100 | 100 | 100 | 100 | 100 |
| 358 | 153 | 98.2 | 0.9 | 96.7 | 97.7 | 97.9 | 98.3 | 100 |
| 316 | 276 | 95 | 3.8 | 83.5 | 94.5 | 96 | 97.9 | 99.8 |
| 32022 | 435 | 99.3 | 0.3 | 98.6 | 99.1 | 99.2 | 99.5 | 100 |
| 321 | 3 | 99.9 | 0.1 | 99.8 | 99.8 | 99.9 | 99.9 | 99.9 |
| 337330 | 21 | 98.9 | 0.8 | 98.1 | 98.1 | 98.2 | 99.7 | 100 |
| 54571 | 36 | 91.1 | 4 | 84 | 89.3 | 91 | 92.2 | 99.8 |
| 546 | 7750 | 99.5 | 0.5 | 98.5 | 98.7 | 99.7 | 99.9 | 100 |
| 106654 | 45 | 99.7 | 0.1 | 99.6 | 99.7 | 99.7 | 99.8 | 99.9 |
| 983594 | 3 | 99.9 | 0.1 | 99.9 | 99.9 | 99.9 | 100 | 100 |
| 98360 | 36 | 100 | 0 | 100 | 100 | 100 | 100 | 100 |
| 28110 | 15 | 98 | 1.2 | 96.2 | 97.5 | 97.6 | 98.8 | 100 |
| 163164 | 15 | 100 | 0 | 100 | 100 | 100 | 100 | 100 |

| | | | | | | | | |
|---|---|---|---|---|---|---|---|---|
| 202752 | 3 | 99.9 | 0.1 | 99.9 | 99.9 | 99.9 | 100 | 100 |
| 2027860 | 21 | 99.5 | 0.9 | 98.1 | 98.1 | 100 | 100 | 100 |
| 2027919 | 78 | 99.6 | 0.3 | 99.1 | 99.3 | 99.7 | 99.9 | 100 |
| 239935 | 2415 | 98.8 | 1.4 | 95.4 | 97.7 | 99.6 | 99.7 | 100 |
| 33010 | 15 | 99.1 | 0.4 | 98.6 | 98.8 | 98.8 | 99.5 | 99.7 |
| 2012495 | 6 | 99.8 | 0.1 | 99.7 | 99.7 | 99.8 | 99.9 | 99.9 |
| 2014 | 3 | 99.7 | 0.2 | 99.5 | 99.5 | 99.6 | 99.8 | 99.9 |
| 2016517 | 3 | 100 | 0 | 100 | 100 | 100 | 100 | 100 |
| 93218 | 10 | 99.9 | 0 | 99.9 | 99.9 | 99.9 | 100 | 100 |
| 93220 | 10 | 99.8 | 0.1 | 99.7 | 99.7 | 99.7 | 99.9 | 100 |
| 1502 | 120 | 99.2 | 0.7 | 97.1 | 98.7 | 99.7 | 99.8 | 100 |
| 1499973 | 15 | 100 | 0 | 99.9 | 99.9 | 100 | 100 | 100 |
| 150055 | 10 | 99.8 | 0.1 | 99.7 | 99.7 | 99.8 | 100 | 100 |
| 150056 | 3 | 99.9 | 0.1 | 99.9 | 99.9 | 99.9 | 100 | 100 |
| 231049 | 6 | 97.5 | 1.9 | 96.3 | 96.3 | 96.3 | 98.9 | 100 |
| 29379 | 3 | 99.9 | 0.1 | 99.9 | 99.9 | 99.9 | 100 | 100 |
| 29380 | 6 | 100 | 0 | 99.9 | 100 | 100 | 100 | 100 |
| 29385 | 91 | 99.9 | 0.1 | 99.7 | 99.8 | 99.9 | 99.9 | 100 |
| 293387 | 210 | 99.9 | 0.1 | 99.8 | 99.8 | 99.9 | 99.9 | 100 |
| 996 | 465 | 99.6 | 0.7 | 96.9 | 99.9 | 99.9 | 100 | 100 |
| 29494 | 6 | 99.9 | 0.1 | 99.8 | 99.8 | 99.8 | 99.9 | 99.9 |
| 29495 | 6 | 98.3 | 1.4 | 96.8 | 97 | 98.2 | 99.5 | 99.8 |
| 29570 | 3 | 98.9 | 0.5 | 98.6 | 98.6 | 98.6 | 99 | 99.4 |
| 55601 | 66 | 99.9 | 0.1 | 99.8 | 99.8 | 99.8 | 99.9 | 100 |
| 553814 | 6 | 99.3 | 0.4 | 99 | 99 | 99.2 | 99.3 | 100 |
| 554 | 3 | 99.6 | 0.2 | 99.5 | 99.5 | 99.5 | 99.7 | 99.8 |
| 67827 | 15 | 99.1 | 0.7 | 98.2 | 98.4 | 98.6 | 99.8 | 100 |

| | | | | | | | | |
|---|---|---|---|---|---|---|---|---|
| 679895 | 3 | 100 | 0 | 100 | 100 | 100 | 100 | 100 |
| 680 | 21 | 98.3 | 1.3 | 97 | 97.4 | 97.4 | 99.8 | 100 |
| 454 | 3 | 99.4 | 0.1 | 99.4 | 99.4 | 99.4 | 99.5 | 99.5 |
| 53344 | 3 | 99.8 | 0.2 | 99.7 | 99.7 | 99.7 | 99.8 | 100 |
| 53345 | 6 | 99.9 | 0.1 | 99.8 | 99.8 | 99.8 | 99.9 | 100 |
| 53346 | 6 | 99.8 | 0.2 | 99.6 | 99.7 | 99.8 | 99.9 | 100 |
| 202751 | 3 | 100 | 0 | 100 | 100 | 100 | 100 | 100 |
| 28132 | 28 | 99 | 0.5 | 98.4 | 98.6 | 98.8 | 99.5 | 99.7 |
| 1302 | 66 | 99.4 | 0.2 | 99.3 | 99.3 | 99.4 | 99.4 | 100 |
| 1305 | 15 | 99 | 0.1 | 98.8 | 98.9 | 99 | 99.2 | 99.3 |
| 34073 | 3 | 94.7 | 1.2 | 93.8 | 94 | 94.3 | 95.2 | 96 |
| 34085 | 120 | 99.3 | 0.4 | 98.8 | 99.1 | 99.2 | 99.7 | 100 |
| 274 | 105 | 99.1 | 0.7 | 97.4 | 99.1 | 99.3 | 99.7 | 100 |
| 238 | 10 | 99.9 | 0.1 | 99.8 | 99.8 | 100 | 100 | 100 |
| 1392 | 1081 | 97.2 | 2.4 | 92.2 | 95.3 | 96.3 | 99.7 | 100 |
| 2585117 | 3 | 99.7 | 0.2 | 99.6 | 99.6 | 99.6 | 99.8 | 100 |
| 672 | 153 | 99.7 | 0.2 | 99.3 | 99.6 | 99.7 | 99.8 | 100 |
| 85698 | 120 | 98.9 | 1 | 96.7 | 98.7 | 99.3 | 99.7 | 100 |
| 364106 | 3 | 100 | 0 | 100 | 100 | 100 | 100 | 100 |
| 364410 | 28 | 99 | 0.5 | 98.2 | 98.8 | 98.9 | 99.2 | 100 |
| 36809 | 435 | 99.7 | 0.2 | 99.4 | 99.5 | 99.7 | 99.9 | 100 |
| 371601 | 10 | 99.5 | 0.2 | 99.1 | 99.4 | 99.6 | 99.6 | 99.8 |
| 373 | 6 | 98.9 | 0.4 | 98.5 | 98.6 | 99 | 99.2 | 99.5 |
| 42897 | 3 | 100 | 0 | 100 | 100 | 100 | 100 | 100 |
| 56 | 3 | 93.2 | 0.8 | 92.5 | 92.8 | 93.2 | 93.6 | 94 |
| 90370 | 6216 | 100 | 0.2 | 98.7 | 100 | 100 | 100 | 100 |
| 1481663 | 6 | 99.7 | 0.1 | 99.6 | 99.6 | 99.7 | 99.8 | 99.9 |

| | | | | | | | | |
|---|---|---|---|---|---|---|---|---|
| 148814 | 3 | 99.6 | 0.1 | 99.6 | 99.6 | 99.6 | 99.7 | 99.7 |
| 148942 | 15 | 99.7 | 0.4 | 99.2 | 99.2 | 100 | 100 | 100 |
| 1491 | 630 | 93.2 | 8.4 | 75.3 | 94.7 | 96.5 | 98.9 | 100 |
| 180957 | 136 | 99.9 | 0.1 | 99.6 | 99.8 | 99.9 | 99.9 | 100 |
| 1812935 | 190 | 99.8 | 0.2 | 98.9 | 99.9 | 99.9 | 99.9 | 100 |
| 1888 | 6 | 93.9 | 6.2 | 88 | 88.4 | 94 | 99.5 | 99.9 |
| 74426 | 6 | 99.5 | 0.2 | 99.3 | 99.4 | 99.5 | 99.5 | 100 |
| 1292 | 28 | 99.7 | 0.2 | 99.3 | 99.5 | 99.8 | 99.9 | 99.9 |
| 1288 | 15 | 99.8 | 0.1 | 99.6 | 99.7 | 99.7 | 99.9 | 100 |
| 1290 | 10 | 99.8 | 0.1 | 99.6 | 99.7 | 99.8 | 99.8 | 99.9 |
| 91890 | 10 | 99.8 | 0.1 | 99.7 | 99.7 | 99.8 | 99.9 | 100 |
| 91891 | 120 | 99.6 | 0.2 | 99.4 | 99.4 | 99.5 | 99.6 | 100 |
| 1874630 | 3 | 99.8 | 0.1 | 99.8 | 99.8 | 99.8 | 99.8 | 99.9 |
| 2605619 | 45 | 100 | 0 | 100 | 100 | 100 | 100 | 100 |
| 1898207 | 10 | 67.4 | 16.9 | 55.3 | 56.8 | 57.5 | 81.6 | 94.6 |
| 1901 | 91 | 98.6 | 1.9 | 96.1 | 96.2 | 100 | 100 | 100 |
| 881260 | 28 | 99.9 | 0 | 99.9 | 99.9 | 99.9 | 100 | 100 |
| 302911 | 21 | 100 | 0 | 99.9 | 99.9 | 100 | 100 | 100 |
| 28901 | 3916 | 99.6 | 0.7 | 96.1 | 99.4 | 99.9 | 99.9 | 100 |
| 50719 | 10 | 99.8 | 0.1 | 99.7 | 99.8 | 99.8 | 99.9 | 99.9 |
| 2686077 | 3 | 99.3 | 0.6 | 99 | 99 | 99 | 99.5 | 100 |
| 756892 | 171 | 99.5 | 0.2 | 99 | 99.4 | 99.6 | 99.7 | 100 |
| 75985 | 2775 | 99.9 | 0.1 | 99.8 | 99.9 | 99.9 | 100 | 100 |
| 1328 | 21 | 98.5 | 0.5 | 97.9 | 98.2 | 98.5 | 98.8 | 99.7 |
| 1329 | 15 | 99.6 | 0.2 | 99.4 | 99.4 | 99.7 | 99.7 | 100 |
| 1346 | 28 | 100 | 0 | 99.9 | 99.9 | 100 | 100 | 100 |
| 580165 | 91 | 99.6 | 0.4 | 99.1 | 99.2 | 100 | 100 | 100 |

| | | | | | | | | |
|---|---|---|---|---|---|---|---|---|
| 58095 | 66 | 100 | 0 | 100 | 100 | 100 | 100 | 100 |
| 13689 | 10 | 99.9 | 0.1 | 99.8 | 99.8 | 99.8 | 100 | 100 |
| 13690 | 3 | 99.5 | 0.1 | 99.5 | 99.5 | 99.5 | 99.5 | 99.6 |
| 1307 | 1830 | 99.4 | 0.3 | 98.8 | 99.2 | 99.3 | 99.5 | 100 |
| 1404 | 136 | 98 | 4.8 | 84.9 | 99.7 | 99.8 | 99.8 | 100 |
| 1404367 | 6 | 99.6 | 0.2 | 99.4 | 99.5 | 99.6 | 99.7 | 99.8 |
| 1405 | 780 | 96.5 | 2.7 | 90.9 | 94 | 96.1 | 99.3 | 100 |
| 1406 | 45 | 95.8 | 2.7 | 92.5 | 93 | 95.7 | 98.2 | 99.9 |
| 1421 | 15 | 95.5 | 3.1 | 90.6 | 93 | 96.1 | 97.5 | 100 |
| 1423 | 6670 | 99.5 | 1 | 95.9 | 99.8 | 99.8 | 99.9 | 100 |
| 955 | 21 | 95 | 4.9 | 89.9 | 89.9 | 98.9 | 99.8 | 100 |
| 96345 | 91 | 96.3 | 5.8 | 86.3 | 86.7 | 99.7 | 99.9 | 100 |
| 964 | 3 | 99.7 | 0.2 | 99.6 | 99.7 | 99.7 | 99.8 | 99.9 |
| 1588 | 3 | 99.6 | 0.2 | 99.5 | 99.5 | 99.6 | 99.7 | 99.8 |
| 158822 | 6 | 99.4 | 0.3 | 99.1 | 99.2 | 99.4 | 99.5 | 99.8 |
| 41170 | 3 | 99.7 | 0.2 | 99.6 | 99.7 | 99.7 | 99.8 | 99.9 |
| 412384 | 3 | 99.8 | 0.1 | 99.7 | 99.7 | 99.7 | 99.8 | 99.9 |
| 47880 | 3 | 99.9 | 0.1 | 99.9 | 99.9 | 99.9 | 100 | 100 |
| 47883 | 15 | 99.6 | 0.2 | 99.3 | 99.4 | 99.7 | 99.8 | 99.9 |
| 1590 | 8256 | 98.9 | 1.1 | 92.3 | 98.1 | 99 | 99.8 | 100 |
| 1580596 | 21 | 99.3 | 0.4 | 98.8 | 99 | 99 | 99.5 | 100 |
| 1582 | 3 | 99.6 | 0.1 | 99.5 | 99.5 | 99.6 | 99.6 | 99.6 |
| 134875 | 10 | 99.7 | 0.2 | 99.4 | 99.5 | 99.8 | 99.9 | 100 |
| 1349 | 3 | 99.8 | 0 | 99.8 | 99.8 | 99.8 | 99.8 | 99.8 |
| 366649 | 28 | 100 | 0 | 100 | 100 | 100 | 100 | 100 |
| 357276 | 55 | 99.2 | 1.3 | 96.3 | 99.8 | 99.8 | 99.9 | 100 |
| 357441 | 15 | 99.9 | 0.1 | 99.8 | 99.8 | 99.9 | 99.9 | 100 |

| | | | | | | | | |
|---|---|---|---|---|---|---|---|---|
| 35755 | 6 | 99.9 | 0.1 | 99.9 | 99.9 | 99.9 | 100 | 100 |
| 72361 | 3 | 97.2 | 2.3 | 95.9 | 95.9 | 95.9 | 97.9 | 99.9 |
| 1422 | 3 | 99.3 | 0.3 | 99.1 | 99.1 | 99.1 | 99.4 | 99.7 |
| 202956 | 10 | 99.8 | 0.1 | 99.7 | 99.7 | 99.7 | 99.8 | 100 |
| 108619 | 36 | 100 | 0 | 99.9 | 99.9 | 100 | 100 | 100 |
| 1081631 | 3 | 99.4 | 0.1 | 99.3 | 99.3 | 99.4 | 99.5 | 99.5 |
| 630 | 21 | 99.7 | 0.2 | 99.4 | 99.5 | 99.6 | 99.9 | 100 |
| 2096 | 15 | 99.4 | 0.2 | 99.2 | 99.3 | 99.3 | 99.4 | 100 |
| 33940 | 3 | 99.9 | 0.1 | 99.9 | 99.9 | 99.9 | 100 | 100 |
| 33941 | 3 | 99.6 | 0.2 | 99.4 | 99.5 | 99.6 | 99.7 | 99.8 |
| 33945 | 3 | 97.9 | 1.7 | 96.9 | 96.9 | 96.9 | 98.3 | 99.8 |
| 33959 | 45 | 99.6 | 0.1 | 99.4 | 99.5 | 99.6 | 99.7 | 100 |
| 33964 | 10 | 99.7 | 0.1 | 99.6 | 99.6 | 99.8 | 99.8 | 100 |
| 820 | 6 | 99.9 | 0 | 99.9 | 99.9 | 99.9 | 99.9 | 99.9 |
| 821 | 3 | 99.9 | 0.1 | 99.8 | 99.8 | 99.9 | 99.9 | 99.9 |
| 823 | 36 | 99.7 | 0.2 | 99.3 | 99.6 | 99.7 | 99.8 | 99.9 |
| 82633 | 3 | 95.8 | 1.1 | 95 | 95.2 | 95.4 | 96.2 | 97 |
| 82689 | 3 | 100 | 0 | 100 | 100 | 100 | 100 | 100 |
| 305 | 4005 | 99.1 | 1 | 96.2 | 97.8 | 99.9 | 99.9 | 100 |
| 644 | 171 | 99.5 | 0.6 | 97.5 | 99.5 | 99.8 | 99.8 | 100 |
| 43657 | 3 | 100 | 0 | 100 | 100 | 100 | 100 | 100 |
| 43674 | 3 | 100 | 0 | 100 | 100 | 100 | 100 | 100 |
| 43675 | 3 | 99.4 | 0.1 | 99.3 | 99.3 | 99.4 | 99.4 | 99.4 |
| 43765 | 10 | 99.6 | 0.2 | 99.4 | 99.4 | 99.5 | 99.7 | 100 |
| 43767 | 66 | 99.9 | 0 | 99.9 | 99.9 | 99.9 | 99.9 | 100 |
| 29394 | 105 | 99.6 | 0.1 | 99.5 | 99.6 | 99.6 | 99.6 | 100 |
| 29397 | 10 | 99.8 | 0.1 | 99.7 | 99.7 | 99.9 | 99.9 | 100 |

| | | | | | | | | |
|---|---|---|---|---|---|---|---|---|
| 29430 | 78 | 99.8 | 0.2 | 99.3 | 99.8 | 99.9 | 99.9 | 100 |
| 1509 | 15 | 97 | 1.5 | 95.2 | 96 | 96.4 | 98.5 | 100 |
| 32009 | 3 | 97.7 | 0.4 | 97.4 | 97.6 | 97.7 | 97.9 | 98.1 |
| 32019 | 6 | 99.8 | 0.2 | 99.5 | 99.6 | 99.8 | 99.9 | 100 |
| 32021 | 6 | 99.6 | 0.1 | 99.5 | 99.5 | 99.5 | 99.6 | 99.7 |
| 273384 | 15 | 99.6 | 0.3 | 99.1 | 99.2 | 99.8 | 99.9 | 99.9 |
| 1682 | 45 | 99.2 | 0.4 | 98.7 | 98.9 | 99.3 | 99.5 | 100 |
| 615 | 1770 | 99.6 | 0.7 | 95.8 | 99.6 | 99.7 | 99.7 | 100 |
| 648 | 153 | 99.7 | 0.1 | 99.4 | 99.6 | 99.7 | 99.8 | 100 |
| 666 | 1953 | 99.7 | 0.2 | 99 | 99.6 | 99.7 | 99.8 | 100 |
| 542 | 6 | 100 | 0 | 100 | 100 | 100 | 100 | 100 |
| 54291 | 105 | 99.2 | 1 | 96.8 | 98.1 | 99.9 | 100 | 100 |
| 54388 | 6 | 100 | 0 | 100 | 100 | 100 | 100 | 100 |
| 545 | 36 | 99.6 | 0.7 | 98.3 | 99.9 | 99.9 | 100 | 100 |
| 440524 | 820 | 100 | 0 | 100 | 100 | 100 | 100 | 100 |
| 28188 | 6 | 99.3 | 0.1 | 99.2 | 99.2 | 99.3 | 99.3 | 99.5 |
| 28198 | 10 | 99 | 0.4 | 98.6 | 98.7 | 99 | 99.4 | 99.7 |
| 129338 | 3 | 97.8 | 1.7 | 96.8 | 96.8 | 96.8 | 98.3 | 99.8 |
| 129394 | 78 | 99.9 | 0.1 | 99.8 | 99.8 | 99.9 | 99.9 | 100 |
| 1597 | 435 | 99.7 | 0.3 | 98.9 | 99.6 | 99.8 | 99.8 | 100 |
| 654 | 136 | 99.5 | 0.3 | 98.7 | 99.2 | 99.6 | 99.8 | 99.9 |
| 54736 | 6 | 99.9 | 0.1 | 99.8 | 99.8 | 99.8 | 99.9 | 99.9 |
| 548 | 465 | 98.4 | 1.2 | 95.5 | 98 | 98.1 | 99.9 | 100 |
| 546165 | 21 | 100 | 0 | 100 | 100 | 100 | 100 | 100 |
| 595 | 45 | 99.8 | 0.4 | 99 | 100 | 100 | 100 | 100 |
| 58712 | 105 | 100 | 0 | 100 | 100 | 100 | 100 | 100 |
| 587753 | 55 | 99.1 | 1.2 | 96.6 | 99.5 | 99.6 | 99.7 | 100 |

| | | | | | | | | |
|---|---|---|---|---|---|---|---|---|
| 111844 | 3 | 99.6 | 0.1 | 99.5 | 99.5 | 99.6 | 99.6 | 99.6 |
| 386 | 21 | 97.6 | 1.9 | 94.4 | 95.4 | 98.3 | 98.5 | 100 |
| 38323 | 10 | 99.9 | 0.1 | 99.7 | 99.8 | 99.9 | 99.9 | 100 |
| 108980 | 3 | 98.9 | 0.6 | 98.4 | 98.5 | 98.6 | 99.1 | 99.6 |
| 108981 | 10 | 99.4 | 0.1 | 99.1 | 99.3 | 99.4 | 99.4 | 99.6 |
| 2754056 | 3 | 99.5 | 0.3 | 99.3 | 99.4 | 99.5 | 99.7 | 99.8 |
| 1765 | 28 | 99.6 | 0.5 | 98.5 | 99.3 | 99.9 | 100 | 100 |
| 2066070 | 3 | 100 | 0 | 100 | 100 | 100 | 100 | 100 |
| 549 | 36 | 99.7 | 0.3 | 99.2 | 99.8 | 99.9 | 99.9 | 100 |
| 664683 | 3 | 99.8 | 0.1 | 99.8 | 99.8 | 99.8 | 99.8 | 99.9 |
| 1639 | 17205 | 99.7 | 0.2 | 98.9 | 99.6 | 99.7 | 99.9 | 100 |
| 35814 | 1953 | 100 | 0 | 100 | 100 | 100 | 100 | 100 |
| 29486 | 15 | 99.9 | 0.1 | 99.7 | 99.7 | 99.9 | 100 | 100 |
| 29489 | 6 | 99.7 | 0.1 | 99.6 | 99.7 | 99.7 | 99.8 | 99.8 |
| 29491 | 10 | 99.9 | 0.1 | 99.9 | 99.9 | 99.9 | 100 | 100 |
| 1304 | 78 | 99.6 | 0.1 | 99.4 | 99.6 | 99.6 | 99.7 | 100 |
| 69966 | 15 | 99.6 | 0.3 | 99.2 | 99.2 | 99.7 | 99.8 | 99.8 |
| 552 | 36 | 98.4 | 1.4 | 97.2 | 97.2 | 97.2 | 100 | 100 |
| 55212 | 3 | 96.5 | 3 | 94.8 | 94.8 | 94.8 | 97.4 | 100 |
| 553 | 28 | 99.8 | 0.1 | 99.6 | 99.8 | 99.9 | 99.9 | 100 |
| 1262449 | 3 | 100 | 0 | 100 | 100 | 100 | 100 | 100 |
| 1308 | 1953 | 99.7 | 0.3 | 98.9 | 99.6 | 99.7 | 99.8 | 100 |
| 1463165 | 1326 | 99.9 | 0.1 | 99.3 | 99.8 | 99.9 | 100 | 100 |
| 1313 | 1891 | 99.6 | 0.2 | 98.9 | 99.5 | 99.6 | 99.7 | 100 |
| 345073 | 3 | 100 | 0 | 100 | 100 | 100 | 100 | 100 |
| 346 | 10 | 99.5 | 0.2 | 99.2 | 99.3 | 99.5 | 99.6 | 100 |
| 347 | 10 | 99.2 | 0.5 | 98.7 | 98.8 | 99 | 99.5 | 99.9 |

| | | | | | | | | |
|---|---|---|---|---|---|---|---|---|
| 359 | 6 | 94.4 | 6 | 88.9 | 88.9 | 94.4 | 99.9 | 99.9 |
| 38289 | 3 | 87.5 | 10.6 | 81.3 | 81.3 | 81.4 | 90.6 | 99.7 |
| 38290 | 3 | 99.7 | 0 | 99.7 | 99.7 | 99.7 | 99.7 | 99.7 |
| 38294 | 3 | 100 | 0 | 100 | 100 | 100 | 100 | 100 |
| 490 | 3 | 97.3 | 1.1 | 96.5 | 96.7 | 96.8 | 97.7 | 98.6 |
| 492670 | 11175 | 99.7 | 0.3 | 96.5 | 99.7 | 99.8 | 99.9 | 100 |
| 269673 | 6 | 99.9 | 0.1 | 99.9 | 99.9 | 100 | 100 | 100 |
| 714 | 21 | 99.2 | 0.4 | 98.8 | 98.9 | 99 | 99.1 | 99.9 |
| 715 | 3 | 99.2 | 0.6 | 98.9 | 98.9 | 98.9 | 99.4 | 99.9 |
| 208224 | 21 | 98.8 | 1.9 | 95.8 | 95.8 | 99.9 | 100 | 100 |
| 208479 | 3 | 99.8 | 0.2 | 99.7 | 99.7 | 99.7 | 99.8 | 100 |
| 1749 | 6 | 99.7 | 0.1 | 99.7 | 99.7 | 99.7 | 99.8 | 99.8 |
| 1750 | 10 | 98.1 | 1.3 | 96.9 | 97.1 | 97.2 | 99.5 | 100 |
| 1751046 | 3 | 100 | 0 | 100 | 100 | 100 | 100 | 100 |
| 317 | 36 | 98.5 | 2.1 | 94.6 | 99.4 | 99.4 | 99.8 | 100 |
| 29388 | 6 | 99.7 | 0.1 | 99.6 | 99.6 | 99.7 | 99.8 | 99.9 |
| 29391 | 3 | 99.7 | 0.2 | 99.6 | 99.6 | 99.6 | 99.8 | 100 |
| 286783 | 153 | 100 | 0 | 100 | 100 | 100 | 100 | 100 |
| 287 | 44850 | 99.7 | 0.4 | 95.2 | 99.7 | 99.9 | 99.9 | 100 |
| 143221 | 15 | 100 | 0 | 100 | 100 | 100 | 100 | 100 |
| 143387 | 3 | 99.7 | 0.1 | 99.7 | 99.7 | 99.7 | 99.8 | 99.8 |
| 469008 | 6 | 100 | 0 | 100 | 100 | 100 | 100 | 100 |
| 2104 | 253 | 99.9 | 0.1 | 99.5 | 99.8 | 99.8 | 100 | 100 |
| 416213 | 6 | 99.9 | 0.1 | 99.9 | 99.9 | 100 | 100 | 100 |
| 28025 | 28 | 99.8 | 0.3 | 99.5 | 99.5 | 100 | 100 | 100 |
| 645 | 15 | 99.6 | 0.3 | 99.3 | 99.4 | 99.5 | 99.7 | 100 |
| 673 | 6 | 99.9 | 0 | 99.9 | 99.9 | 99.9 | 99.9 | 99.9 |

| | | | | | | | | |
|---|---|---|---|---|---|---|---|---|
| 315405 | 3 | 99.4 | 0.4 | 99.2 | 99.2 | 99.2 | 99.6 | 99.9 |
| 1812934 | 6 | 99.7 | 0.3 | 99.4 | 99.4 | 99.7 | 99.9 | 100 |
| 1038927 | 3 | 100 | 0 | 100 | 100 | 100 | 100 | 100 |
| 104087 | 3 | 96 | 2 | 94.7 | 94.8 | 95 | 96.7 | 98.3 |
| 83555 | 45 | 99.3 | 0.5 | 98.6 | 99 | 99.3 | 99.7 | 100 |
| 83558 | 21 | 100 | 0 | 100 | 100 | 100 | 100 | 100 |
| 83559 | 28 | 99.1 | 0.6 | 98.5 | 98.6 | 98.6 | 99.7 | 99.9 |
| 83560 | 3 | 100 | 0 | 100 | 100 | 100 | 100 | 100 |
| 631 | 6 | 100 | 0 | 100 | 100 | 100 | 100 | 100 |
| 632 | 378 | 100 | 0.1 | 99.5 | 100 | 100 | 100 | 100 |
| 1855823 | 21 | 99.7 | 0.1 | 99.4 | 99.5 | 99.7 | 99.8 | 99.9 |
| 1599 | 153 | 99.7 | 0.2 | 99.1 | 99.7 | 99.8 | 99.9 | 100 |
| 1598 | 136 | 99.2 | 0.6 | 98.1 | 98.4 | 99.5 | 99.6 | 100 |
| 294 | 190 | 96.5 | 2.1 | 88.6 | 96.2 | 96.4 | 97.8 | 100 |
| 565 | 3 | 100 | 0.1 | 99.9 | 100 | 100 | 100 | 100 |
| 564 | 990 | 99.9 | 0.1 | 99.7 | 99.9 | 99.9 | 99.9 | 100 |
| 36855 | 15 | 100 | 0 | 100 | 100 | 100 | 100 | 100 |
| 47917 | 21 | 99.8 | 0.1 | 99.7 | 99.8 | 99.8 | 99.9 | 100 |
| 29347 | 3 | 99.4 | 0.4 | 99.2 | 99.2 | 99.2 | 99.6 | 99.9 |
| 337 | 36 | 98.5 | 2.4 | 93.9 | 99.6 | 99.8 | 99.9 | 100 |
| 33889 | 10 | 98.3 | 2 | 96 | 96 | 99.7 | 99.8 | 100 |
| 1393 | 3 | 98.8 | 0.4 | 98.6 | 98.6 | 98.6 | 98.9 | 99.3 |
| 48664 | 15 | 99.3 | 0.9 | 98.1 | 98.1 | 99.9 | 100 | 100 |
| 240427 | 3 | 99.8 | 0.1 | 99.7 | 99.7 | 99.7 | 99.8 | 99.9 |
| 587851 | 55 | 98.9 | 1.3 | 95.4 | 98.8 | 99.9 | 99.9 | 100 |
| 588 | 45 | 100 | 0 | 99.9 | 99.9 | 100 | 100 | 100 |
| 588858 | 10 | 100 | 0 | 100 | 100 | 100 | 100 | 100 |

| | | | | | | | | |
|---|---|---|---|---|---|---|---|---|
| 588932 | 3 | 98.2 | 1.1 | 97.6 | 97.6 | 97.6 | 98.5 | 99.5 |
| 648995 | 3 | 99.8 | 0 | 99.8 | 99.8 | 99.8 | 99.8 | 99.8 |
| 1571 | 6 | 98.1 | 0.5 | 97.5 | 97.7 | 98 | 98.4 | 98.9 |
| 1600 | 6 | 99.7 | 0.1 | 99.6 | 99.6 | 99.7 | 99.7 | 99.8 |
| 157687 | 6 | 98.9 | 0.4 | 98.4 | 98.6 | 98.8 | 99.1 | 99.4 |
| 157782 | 3 | 97.7 | 1.9 | 96.6 | 96.6 | 96.6 | 98.2 | 99.9 |
| 473426 | 3 | 99.6 | 0.2 | 99.5 | 99.5 | 99.5 | 99.7 | 99.8 |
| 47466 | 10 | 97.8 | 0.9 | 96.8 | 97.4 | 97.6 | 97.9 | 100 |
| 47678 | 6 | 100 | 0.1 | 99.9 | 99.9 | 100 | 100 | 100 |
| 47714 | 6 | 99.5 | 0.2 | 99.3 | 99.3 | 99.5 | 99.6 | 99.7 |
| 147375 | 10 | 95 | 3 | 91 | 93.4 | 94.8 | 97.3 | 100 |
| 487838 | 15 | 100 | 0 | 100 | 100 | 100 | 100 | 100 |
| 28031 | 3 | 92.4 | 2.3 | 90 | 91.3 | 92.7 | 93.7 | 94.6 |
| 488 | 3 | 97.6 | 1.8 | 96.5 | 96.5 | 96.6 | 98.2 | 99.7 |
| 488446 | 3 | 99.9 | 0.1 | 99.9 | 99.9 | 99.9 | 100 | 100 |
| 488447 | 10 | 98.3 | 1.3 | 97.2 | 97.3 | 97.4 | 99.8 | 99.8 |
| 1314 | 23220 | 99.6 | 0.2 | 96.3 | 99.6 | 99.6 | 99.7 | 100 |
| 611 | 496 | 100 | 0 | 100 | 100 | 100 | 100 | 100 |
| 113557 | 36 | 99.9 | 0.1 | 99.8 | 99.8 | 99.9 | 99.9 | 100 |
| 1138383 | 36 | 99.7 | 0.3 | 99 | 99.7 | 99.8 | 99.9 | 100 |
| 1110693 | 6 | 100 | 0 | 100 | 100 | 100 | 100 | 100 |
| 1660067 | 3 | 99.6 | 0 | 99.6 | 99.6 | 99.6 | 99.6 | 99.6 |
| 126385 | 55 | 99.6 | 0.5 | 98.8 | 99.1 | 99.9 | 99.9 | 100 |
| 1363 | 45 | 99.1 | 0.9 | 97.4 | 99.1 | 99.4 | 99.8 | 100 |
| 1366 | 15 | 99.8 | 0.1 | 99.7 | 99.7 | 99.8 | 99.8 | 99.9 |
| 1282 | 2346 | 99.8 | 0.2 | 99.4 | 99.8 | 99.9 | 99.9 | 100 |
| 1617964 | 3 | 100 | 0 | 100 | 100 | 100 | 100 | 100 |

| | | | | | | | | |
|---|---|---|---|---|---|---|---|---|
| 575 | 10 | 99.9 | 0.1 | 99.9 | 99.9 | 99.9 | 100 | 100 |
| 576610 | 15 | 93.9 | 1.2 | 92.5 | 93.1 | 93.7 | 94.8 | 96.6 |
| 587 | 300 | 99 | 0.9 | 96.9 | 98.3 | 99.3 | 99.7 | 100 |
| 104609 | 10 | 100 | 0 | 100 | 100 | 100 | 100 | 100 |
| 1283 | 136 | 99.1 | 0.9 | 95.9 | 99.2 | 99.4 | 99.8 | 100 |
| 611301 | 630 | 100 | 0 | 100 | 100 | 100 | 100 | 100 |
| 614 | 15 | 99.9 | 0 | 99.9 | 99.9 | 99.9 | 100 | 100 |
| 61645 | 171 | 99.8 | 0.1 | 99.4 | 99.8 | 99.8 | 99.9 | 100 |
| 623 | 190 | 99.9 | 0.1 | 99.7 | 99.9 | 99.9 | 100 | 100 |
| 29461 | 21 | 99.9 | 0.1 | 99.8 | 99.9 | 99.9 | 100 | 100 |
| 29466 | 6 | 99.5 | 0.3 | 99.3 | 99.3 | 99.5 | 99.5 | 100 |
| 29471 | 10 | 100 | 0 | 100 | 100 | 100 | 100 | 100 |
| 817 | 136 | 99.4 | 0.4 | 98.9 | 99 | 99.1 | 99.9 | 100 |
| 818 | 21 | 99.9 | 0 | 99.8 | 99.9 | 99.9 | 99.9 | 100 |
| 85404 | 3 | 99.1 | 0.1 | 99 | 99 | 99.1 | 99.2 | 99.2 |
| 85581 | 6 | 99 | 1 | 98 | 98.1 | 99 | 99.9 | 100 |
| 1496 | 3486 | 99.6 | 0.4 | 98.9 | 99.3 | 99.7 | 99.9 | 100 |
| 246196 | 3 | 99.4 | 0.2 | 99.1 | 99.3 | 99.5 | 99.5 | 99.5 |
| 224308 | 6 | 100 | 0 | 100 | 100 | 100 | 100 | 100 |
| 2371 | 15 | 99.8 | 0.1 | 99.7 | 99.7 | 99.7 | 99.8 | 100 |
| 122 | 3 | 98.3 | 1.4 | 97.4 | 97.5 | 97.6 | 98.8 | 99.9 |
| 195709 | 6 | 99.7 | 0.2 | 99.6 | 99.6 | 99.6 | 99.8 | 100 |
| 196 | 21 | 99.9 | 0.1 | 99.8 | 99.8 | 99.9 | 100 | 100 |
| 1397 | 3 | 97.5 | 1.5 | 96.4 | 96.7 | 97 | 98.1 | 99.2 |
| 1398 | 45 | 99.3 | 0.5 | 97.8 | 98.9 | 99.4 | 99.7 | 100 |
| 149539 | 2211 | 100 | 0.2 | 99 | 100 | 100 | 100 | 100 |
| 57975 | 55 | 98.7 | 2.6 | 93.1 | 99.7 | 99.9 | 99.9 | 100 |

| | | | | | | | | |
|---|---|---|---|---|---|---|---|---|
| 24 | 21 | 98.2 | 1.2 | 96.4 | 97.1 | 97.9 | 99.9 | 100 |
| 232537 | 3 | 99.8 | 0.1 | 99.7 | 99.8 | 99.8 | 99.8 | 99.8 |
| 152331 | 3 | 98.8 | 1 | 98.2 | 98.2 | 98.2 | 99.1 | 100 |
| 1465 | 3 | 98.8 | 1 | 98.2 | 98.2 | 98.2 | 99.1 | 99.9 |
| 146919 | 15 | 99.6 | 0.2 | 99.3 | 99.5 | 99.5 | 99.8 | 99.9 |
| 1338 | 10 | 99.1 | 0.3 | 98.9 | 99 | 99 | 99 | 100 |
| 624 | 561 | 100 | 0 | 99.8 | 100 | 100 | 100 | 100 |
| 119219 | 3 | 99.9 | 0.1 | 99.8 | 99.8 | 99.8 | 99.9 | 100 |
| 585 | 21 | 99.3 | 0.3 | 99 | 99 | 99.2 | 99.3 | 100 |
| 312306 | 3 | 96.5 | 2.9 | 94.8 | 94.8 | 94.8 | 97.3 | 99.8 |
| 1747 | 78 | 99.7 | 0.3 | 98.9 | 99.5 | 99.9 | 100 | 100 |
| 172042 | 6 | 97 | 2.7 | 94.3 | 94.4 | 97.1 | 99.5 | 99.5 |
| 172045 | 3 | 99.4 | 0.2 | 99.3 | 99.3 | 99.3 | 99.5 | 99.7 |
| 728 | 45 | 99.8 | 0.4 | 99 | 100 | 100 | 100 | 100 |
| 61646 | 6 | 99 | 1 | 98.1 | 98.1 | 99 | 99.9 | 100 |
| 61647 | 3 | 98.8 | 1 | 98.2 | 98.2 | 98.3 | 99.1 | 99.9 |
| 123899 | 6 | 99.9 | 0.1 | 99.9 | 99.9 | 99.9 | 100 | 100 |
| 600 | 6 | 100 | 0 | 100 | 100 | 100 | 100 | 100 |
| 60520 | 3 | 98.6 | 0.8 | 97.9 | 98.1 | 98.3 | 98.9 | 99.5 |
| 60550 | 3 | 99 | 0.2 | 98.9 | 98.9 | 98.9 | 99.1 | 99.3 |
| 1639133 | 66 | 99.9 | 0 | 99.8 | 99.9 | 99.9 | 99.9 | 100 |
| 1642 | 6 | 99.8 | 0.1 | 99.8 | 99.8 | 99.8 | 99.8 | 100 |
| 28454 | 6 | 99.6 | 0.3 | 99.3 | 99.4 | 99.4 | 99.8 | 100 |
| 536 | 3 | 99.8 | 0.2 | 99.7 | 99.7 | 99.7 | 99.8 | 100 |
| 473421 | 15 | 100 | 0 | 100 | 100 | 100 | 100 | 100 |
| 264 | 6 | 98.4 | 1 | 97.7 | 97.7 | 97.8 | 99.3 | 99.7 |
| 285 | 3 | 97.7 | 1 | 97.1 | 97.1 | 97.1 | 98 | 98.9 |

| | | | | | | | | |
|---|---|---|---|---|---|---|---|---|
| 521 | 10 | 99.9 | 0.1 | 99.8 | 99.8 | 99.9 | 100 | 100 |
| 1446746 | 3 | 100 | 0 | 100 | 100 | 100 | 100 | 100 |
| 1426 | 3 | 100 | 0 | 100 | 100 | 100 | 100 | 100 |
| 487004 | 6 | 100 | 0 | 100 | 100 | 100 | 100 | 100 |
| 487821 | 6 | 100 | 0.1 | 99.9 | 99.9 | 100 | 100 | 100 |
| 691 | 3 | 99.9 | 0 | 99.9 | 99.9 | 99.9 | 99.9 | 99.9 |
| 69218 | 3 | 99.9 | 0.1 | 99.9 | 99.9 | 99.9 | 100 | 100 |
| 91892 | 36 | 99.9 | 0.1 | 99.7 | 99.8 | 99.9 | 100 | 100 |
| 90371 | 5253 | 99.9 | 0.2 | 98.5 | 100 | 100 | 100 | 100 |
| 911022 | 435 | 100 | 0 | 100 | 100 | 100 | 100 | 100 |
| 1589 | 6 | 98.8 | 0.9 | 98.1 | 98.2 | 98.2 | 99.4 | 99.9 |
| 159 | 10 | 99.8 | 0.1 | 99.8 | 99.8 | 99.8 | 99.9 | 99.9 |
| 76856 | 3 | 99.6 | 0.1 | 99.6 | 99.6 | 99.6 | 99.7 | 99.7 |
| 76857 | 3 | 99.3 | 0.1 | 99.2 | 99.2 | 99.3 | 99.3 | 99.4 |
| 76859 | 15 | 99.5 | 0.1 | 99.4 | 99.5 | 99.5 | 99.5 | 99.6 |
| 76860 | 3 | 99 | 0.6 | 98.6 | 98.7 | 98.7 | 99.2 | 99.7 |
| 349520 | 3 | 100 | 0 | 100 | 100 | 100 | 100 | 100 |
| 1275 | 3 | 99.8 | 0.1 | 99.7 | 99.7 | 99.7 | 99.8 | 99.9 |
| 1748 | 6 | 99.8 | 0.2 | 99.7 | 99.7 | 99.7 | 99.9 | 100 |
| 1596 | 10 | 99.7 | 0.3 | 99.3 | 99.4 | 99.7 | 99.9 | 100 |
| 938155 | 3 | 99.9 | 0.1 | 99.8 | 99.8 | 99.8 | 99.9 | 100 |
| 941322 | 6 | 100 | 0 | 99.9 | 100 | 100 | 100 | 100 |
| 2103 | 21 | 98.2 | 1.9 | 94.7 | 95.8 | 98.9 | 99.3 | 100 |
| 2100 | 6 | 99.9 | 0.1 | 99.8 | 99.8 | 99.8 | 99.9 | 100 |
| 1408 | 78 | 99.3 | 0.4 | 98.8 | 99 | 99.3 | 99.5 | 99.9 |
| 1358 | 55 | 99.6 | 0.2 | 99.3 | 99.4 | 99.6 | 99.8 | 100 |
| 644356 | 21 | 100 | 0 | 99.9 | 99.9 | 100 | 100 | 100 |

| | | | | | | | | |
|---|---|---|---|---|---|---|---|---|
| 644357 | 28 | 99.9 | 0.1 | 99.8 | 99.9 | 99.9 | 100 | 100 |
| 197700 | 3 | 100 | 0 | 100 | 100 | 100 | 100 | 100 |
| 1978231 | 3 | 68.1 | 7.8 | 63.4 | 63.6 | 63.9 | 70.5 | 77.1 |
| 137591 | 45 | 99.8 | 0.1 | 99.7 | 99.7 | 99.8 | 99.8 | 100 |
| 1353 | 3 | 96.7 | 2.8 | 95.1 | 95.1 | 95.1 | 97.5 | 99.9 |
| 138532 | 10 | 99.8 | 0.1 | 99.7 | 99.7 | 99.8 | 99.8 | 99.9 |
| 59201 | 561 | 99.6 | 0.7 | 96.6 | 99.8 | 99.9 | 99.9 | 100 |
| 1890302 | 10 | 97.4 | 2.8 | 94.1 | 94.2 | 99.6 | 99.6 | 99.6 |
| 119856 | 3 | 98.5 | 1.1 | 97.8 | 97.9 | 98 | 98.9 | 99.8 |
| 101571 | 6 | 95.7 | 2.8 | 93.7 | 93.8 | 94 | 98 | 99.4 |
| 1019 | 6 | 99.3 | 0.1 | 99.3 | 99.3 | 99.3 | 99.4 | 99.4 |
| 71237 | 3 | 99.9 | 0.1 | 99.9 | 99.9 | 99.9 | 100 | 100 |
| 292 | 36 | 97.6 | 2.4 | 92.1 | 97.3 | 98.8 | 99.1 | 100 |
| 96344 | 6 | 99.5 | 0.5 | 99 | 99 | 99.5 | 99.9 | 100 |
| 985002 | 21 | 99.9 | 0.1 | 99.8 | 99.8 | 99.8 | 99.9 | 100 |
| 985762 | 3 | 99.9 | 0.1 | 99.9 | 99.9 | 99.9 | 100 | 100 |
| 59202 | 3 | 99.9 | 0.1 | 99.8 | 99.8 | 99.8 | 99.9 | 100 |
| 1055538 | 55 | 100 | 0 | 100 | 100 | 100 | 100 | 100 |
| 44283 | 3 | 99.7 | 0.2 | 99.6 | 99.6 | 99.6 | 99.8 | 99.9 |
| 1281 | 6 | 99.8 | 0.1 | 99.8 | 99.8 | 99.8 | 99.8 | 100 |
| 58097 | 3 | 100 | 0 | 100 | 100 | 100 | 100 | 100 |
| 596 | 10 | 100 | 0 | 100 | 100 | 100 | 100 | 100 |
| 59814 | 6 | 99.1 | 0.8 | 98.3 | 98.4 | 99.2 | 99.9 | 99.9 |
| 211968 | 36 | 100 | 0 | 100 | 100 | 100 | 100 | 100 |
| 226665 | 6 | 100 | 0 | 100 | 100 | 100 | 100 | 100 |
| 34 | 28 | 98.8 | 0.9 | 97.7 | 97.8 | 98.8 | 99.9 | 100 |
| 340 | 15 | 99.6 | 0.3 | 99.2 | 99.4 | 99.5 | 99.8 | 100 |

| | | | | | | | | |
|---|---|---|---|---|---|---|---|---|
| 340190 | 6 | 100 | 0 | 100 | 100 | 100 | 100 | 100 |
| 34021 | 6 | 100 | 0.1 | 99.9 | 99.9 | 100 | 100 | 100 |
| 1246 | 6 | 99.6 | 0.1 | 99.5 | 99.5 | 99.6 | 99.7 | 99.8 |
| 568703 | 6 | 100 | 0 | 100 | 100 | 100 | 100 | 100 |
| 1176649 | 15 | 99.8 | 0.1 | 99.7 | 99.7 | 99.8 | 100 | 100 |
| 726 | 6 | 99.1 | 0.1 | 98.9 | 99 | 99 | 99.2 | 99.2 |
| 141679 | 3 | 99.9 | 0 | 99.9 | 99.9 | 99.9 | 99.9 | 99.9 |
| 1244531 | 3 | 99.5 | 0.3 | 99.3 | 99.3 | 99.3 | 99.5 | 99.8 |
| 626774 | 45 | 99.9 | 0.2 | 99.5 | 99.9 | 99.9 | 100 | 100 |
| 83333 | 190 | 100 | 0 | 99.9 | 100 | 100 | 100 | 100 |
| 83334 | 6786 | 100 | 0 | 99.9 | 100 | 100 | 100 | 100 |
| 83554 | 15 | 99.8 | 0.2 | 99.6 | 99.6 | 99.7 | 99.9 | 100 |
| 1076 | 10 | 98.4 | 0.6 | 97.6 | 98 | 98.2 | 98.9 | 99.4 |
| 1694 | 6 | 98.6 | 1.6 | 97.1 | 97.1 | 98.5 | 100 | 100 |
| 260678 | 6 | 100 | 0 | 100 | 100 | 100 | 100 | 100 |
| 470934 | 10 | 97.6 | 2.4 | 94.8 | 94.9 | 99.2 | 99.5 | 99.8 |
| 471 | 3 | 99.3 | 0.3 | 99.1 | 99.2 | 99.3 | 99.4 | 99.6 |
| 59204 | 10 | 98.5 | 1.3 | 97.5 | 97.5 | 97.5 | 100 | 100 |
| 593905 | 3 | 99.9 | 0.1 | 99.9 | 99.9 | 99.9 | 100 | 100 |
| 2026186 | 10 | 95.5 | 2.1 | 92.3 | 93.7 | 96 | 96.9 | 99.1 |
